# The lupus susceptibility allele *DRB1*03:01* encodes a disease-driving epitope

**DOI:** 10.1101/2022.05.31.494172

**Authors:** Bruna Miglioranza Scavuzzi, Vincent van Drongelen, Bhavneet Kaur, Jennifer Callahan Fox, Jianhua Liu, Raquel A. Mesquita-Ferrari, J. Michelle Kahlenberg, Evan A. Farkash, Fernando Benavides, Frederick W. Miller, Amr H. Sawalha, Joseph Holoshitz

**Affiliations:** Department of Internal Medicine, University of Michigan, Ann Arbor, MI, 48109, USA; Department of Pathology, University of Michigan, Ann Arbor, MI, 48109, USA; Department of Epigenetics and Molecular Carcinogenesis, MD Anderson Cancer Center, Houston, TX, 77030, USA; Environmental Autoimmunity Group, National Institute of Environmental Health Sciences, Research Triangle Park, NC, 27709, USA; Departments of Pediatrics and Internal Medicine, University of Pittsburgh, Pittsburgh, PA 15224, USA

**Keywords:** Autoimmune disease, Systemic lupus erythematosus, HLA-disease association, Endoplasmic reticulum, Mitochondrial dysfunction, RNA-sequencing, Lupus nephritis, Interferon gamma

## Abstract

The *HLA-DRB1*03:01* allele is a major genetic risk factor in systemic lupus erythematosus (SLE), but the mechanistic basis of the association is unclear. Here we show that in the presence of interferon gamma (IFN-γ), a short *DRB1*03:01*-encoded allelic epitope activates a characteristic lupus transcriptome in mouse and human macrophages. It also triggers a cascade of SLE-associated cellular aberrations, including endoplasmic reticulum stress, unfolded protein response, mitochondrial dysfunction, necroptotic cell death, and production of pro-inflammatory cytokines. Parenteral administration of IFN-γ to naïve *DRB1*03:01* transgenic mice causes increased serum levels of anti-double stranded DNA antibodies, glomerular immune complex deposition and histopathological renal changes that resemble human lupus nephritis. This study provides evidence for a noncanonical, antigen presentation-independent mechanism of HLA-disease association in SLE and could lay new foundations for our understanding of key molecular mechanisms that trigger and propagate this devastating autoimmune disease.

## INTRODUCTION

Systemic lupus erythematosus (SLE) is an autoimmune disease that afflicts millions of individuals worldwide ^1^. The main impediment to finding a cure for SLE is incomplete understanding of its genetic and molecular mechanisms. Salient cellular aberrations observed in SLE include enhanced endoplasmic reticulum (ER) stress, an activated unfolded protein response (UPR), mitochondrial dysfunction and aberrant cell death, associated with a pro-inflammatory state and generation of autoantibodies against nuclear antigens, which are implicated in target tissue damage, such as lupus nephritis (LN) ^1–4^. The molecular mechanism that triggers this cascade of events is largely unknown.

The contributions of genetic factors to SLE risk or disease severity are, likewise, incompletely understood. Susceptibility to the disease has been associated with more than 100 loci ^5,6^, with a particularly strong involvement of human leukocyte antigen (HLA) genes ^7,8^. Meta-analysis of genomic studies has identified *DRB1*03:01* as the single most significant SLE-associated HLA allele in Europeans ^9^. The underlying mechanism, however, is unknown. The prevailing hypotheses concerning HLA-disease association postulate presentation of self or foreign antigens by HLA molecules ^4,8^; however, the antigen presentation hypotheses have not yet been empirically validated in SLE.

Seeking better understanding of the functional role of HLA molecules in SLE, we examined an antigen presentation-independent mechanism, modeled after the ‘MHC Cusp’ theory ^10,11^, which postulates that in addition to presenting antigenic peptides to T cells, major histocompatibility complex (MHC) molecules express signal transducing ligands that, upon binding to non-MHC receptors trigger allele-specific cell activation events. Under certain environmental circumstances and permissive background genes, this activation can provoke aberrant cellular events that facilitate autoimmune diseases. At the focus of the MHC Cusp theory is an α-helical cusp-like conformational motif that is shared by all products of the MHC gene family, irrespective of their primary sequences, and independent of antigen presentation. In HLA-DR molecules, the cusp involves the third allelic hypervariable region (TAHR; residues 65-79) of the DRβ chain. In rheumatoid arthritis (RA) the cusp region that is coded by disease-associated *HLA-DRB1* alleles encompasses a ‘shared epitope’ (SE), which has been shown to act as a ligand that activates *in vitro* and *in vivo* pro-arthritogenic events in mice ^12–17^

Here, we examined whether the TAHR of the DRβ chain coded by the SLE-susceptibility allele *DRB1*03:01* may be directly contributing to SLE pathogenesis. Our findings reveal that reminiscent of SE-coding *DRB1* alleles in RA, the SLE-risk allele *DRB1*03:01* encodes what we designated here as a ‘lupus epitope’ (LE) in the TAHR of the DRβ chain that, in the presence of interferon gamma (IFN-γ), activates signature lupus transcriptomes in mouse and human macrophages, and triggers SLE-characteristic cellular aberrations, including ER stress, UPR, mitochondrial dysfunction, necroptosis and production of pro-inflammatory cytokines. In the presence of IFN-γ, primary mouse bone marrow-derived macrophages (BMDMs) from non-immunized transgenic mice that carry the LE-coding *DRB1*03:01* allele exhibit similar cellular aberrations *ex vivo*, and *in vivo* administration of IFN-γ to those transgenic mice produces an SLE-like disease, including formation of anti-double stranded DNA (dsDNA) antibodies and renal immuno-histopathology changes akin to LN.

## RESULTS

### TAHR epitopes activate allele-specific transcriptomic signatures

Macrophages play a central role in autoimmune diseases, including SLE ^12–14^, and IFN-γ, known for its macrophage activation effects, is a significant factor in human SLE and experimental models of the disease ^15,16^. Accordingly, to explore the transcriptional effects of various allelic epitopes in the HLA-DR cusp region (Supplementary Fig. 1a), we studied mouse (RAW 264.7) and human (THP-1) macrophages in the presence or absence of IFN-γ. Cells were incubated with or without one of the following linear synthetic 15mer TAHR peptides (Supplementary Fig.1b): 65-79*LE, a TAHR (residues 65-79 of the DRβ chain) encoded by the SLE-risk allele *DRB1*03:01*; 65-79*SE, a TAHR encoded by a RA-risk allele *DRB1*\**04:01;* or 65-79*PE, a TAHR epitope containing a 70-DERAA-74 motif which is shared by alleles (*e.g. DRB1*04:02, DRB1*13:01* & *DRB1*13:02*), known to associate with autoimmune disease protection ^17–19^. RNA sequencing (RNA-seq) was performed at 72 h, as previously described ^20^.

Quantitative RT-PCR (qRT-PCR) analysis showed no significant effect on epitope-activated gene expression in the absence of IFN-γ. In its presence, however, substantial increased expression was found for representative macrophage activation gene markers (Supplementary Fig.1c). To characterize TAHR epitopes-driven transcriptional landscapes, and to differentiate between the respective contributions of the different epitopes versus IFN-γ, RNA-seq analyses were performed using two models: 1. Transcriptional modulation by allelic epitopes in the presence of IFN-γ, versus IFN-γ alone (Model A), or 2. Transcriptional modulation by allelic epitopes in the presence of IFN-γ, versus epitope alone (Model B).

Using Model A analysis, unsupervised clustering of upregulated and downregulated differentially expressed genes (DEGs) showed distinct patterns for 65-79*LE, versus 65-79*SE or 65-79*PE (Fig. 1a). As shown in Venn diagrams (Fig. 1b) there were 683 unique DEGs upon exposure of RAW 264.7 macrophages to 65-79*LE (433 overexpressed and 250 underexpressed), and 349 unique DEGs upon exposure to 65-79*SE (173 overexpressed and 176 underexpressed genes). This model identified only 11 unique DEGs that were differentially regulated in cells exposed to 65-79*PE (Fig. 1b). Notable among the most significant group of 65-79*LE-upregulated DEGs were multiple SLE-associated genes, including *Irf4*, *Ccl5*, *Cxcl10*, *Pim1*, *Traf1* amongst others (Fig. 1c and Supplementary table 1). By contrast, 65-79*SE-upregulated DEGs included many RA-relevant genes, including *Tnfrsf9, Relb, Tnaifp3* (Fig. 1c and Supplementary Table 1). Complete lists of all 65-79*LE-and 65-79*SE-modulated DEGs are shown in Supplementary Data File 1.

**Fig. 1:**
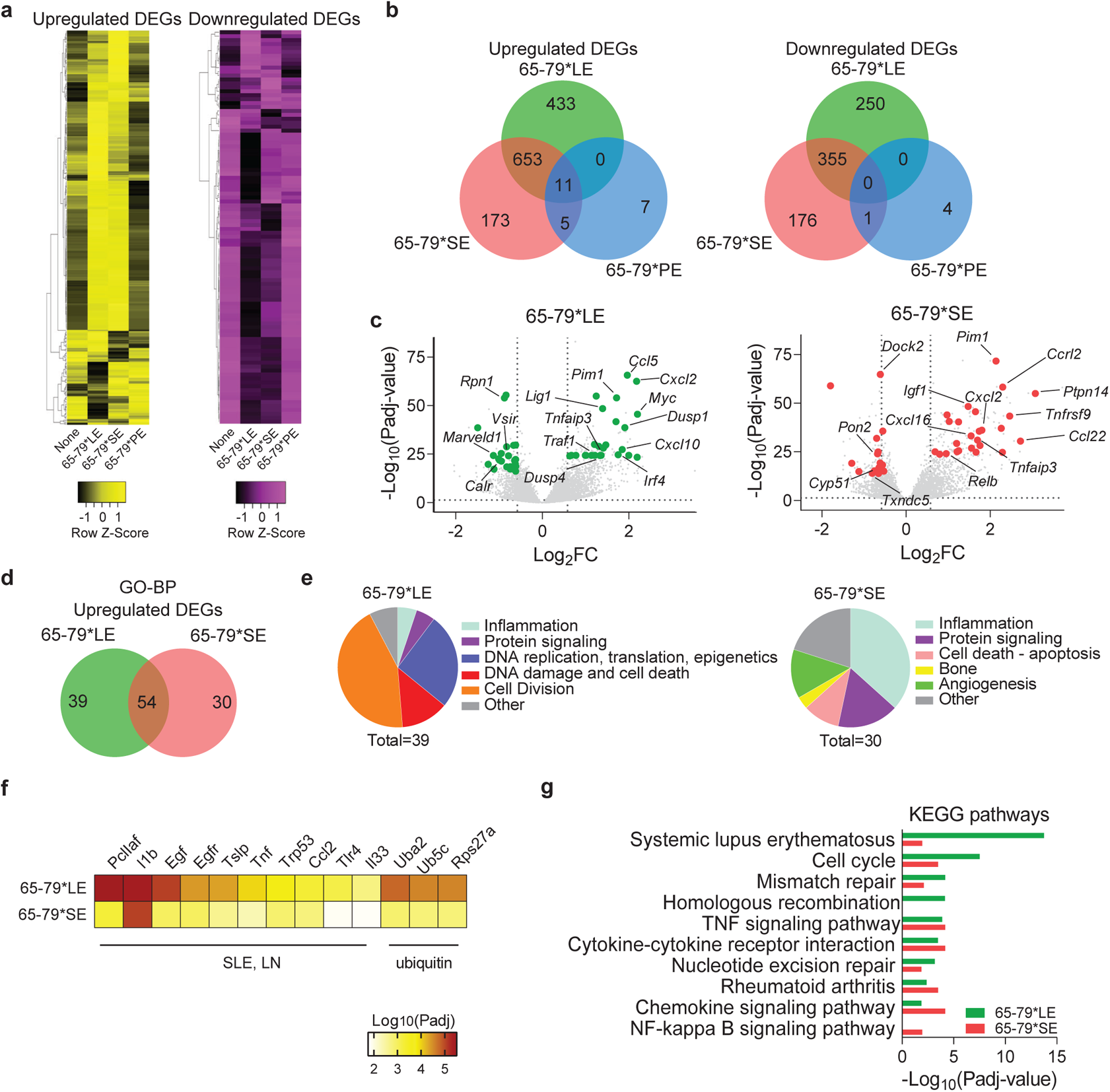
Transcriptional modulation by the LE (Model A: epitope + IFN-γ, versus IFN-γ alone) **a,** Heatmaps showing unsupervised clustering of upregulated (left, yellow) and downregulated (right, purple) DEGs for 65-79*LE, 65-79*SE and 65-79*PE. DEGs were selected based on a > 3-fold change for 65-79*LE. All depicted DEGs had a Padj < 0.05. **b,** Venn diagrams showing comparison of DEGs for 65-79*LE, 65-79*SE and 65-79*PE. **c,** Volcano plots comparing transcriptional modulation by 65-79*LE and 65-79*SE. Green dots denote SLE-relevant genes modulated by 65-79*LE; red dots denote RA-relevant genes modulated by 65-79*SE. **d,** Venn diagram comparing GO-BP terms enrichment among upregulated DEGs for 65-79*LE versus 65-79*SE. **e,** Pie charts showing differential thematic clustering of unique GO-BP term enrichment among 65-79*LE- (left) versus 65-79*SE- (right) upregulated DEGs. **f,** Heatmap of top-ranked SLE- or ubiquitination-associated URs that are predicted to be involved in 65-79*LE transcriptional modulation, as compared to their predicted involvement in 65-79*SE transcriptional modulation. **g,** Notable KEGG pathway enrichment among 65-79*LE-upregulated DEGs, versus 65-79*SE. Data in (**b**-**g**) are based on DEGs with a fold change >1.5 and Padj < 0.05. All data are based on 6 biological replicates in 2 independent experiments. See also Supplementary Tables 1 and 4.

The respective patterns of differential gene expression by 65-79*LE versus 65-79*SE followed distinct, SLE- or RA-relevant functional categories, as determined by Gene Ontology (GO) biological processes (BP) analysis (Figs. 1d and 1e). Additionally, upstream regulator (UR) analysis by iPathwayGuide (iPG) showed distinct patterns between transcriptomes of 65-79*LE- and 65-79*SE-stimulated cells. There were 118 unique URs predicted in RAW 264.7 cells stimulated by 65-79*LE and 6 URs that were uniquely involved in 65-79*SE-stimulated cells. Many of the most significant URs predicted in 65-79*LE-stimulated RAW 264.7 cells are known to play pathogenic roles in SLE or ubiquitination (Fig. 1f, Supplementary Table 4). Intriguingly, Kyoto Encyclopedia of Genes and Genomes (KEGG) pathway analysis (Fig. 1g) identified “systemic lupus erythematosus” as the most significant pathway for upregulated DEGs by 65-79*LE (Padj = 1.79×10^−14^), while “rheumatoid arthritis” (Padj = 3.27×10^−04^) was identified as one of the most significant predicted pathways among DEGs upregulated by 65-79*SE. These data are consistent with the known associations between the LE-coding allele *DRB1*03:01* and SLE ^9^, and between the SE-coding *DRB1*04:01* allele and RA ^21^. Thus, disease-associated allele-specific TAHR epitopes that are antigen presentation-incompetent activate distinct, disease-relevant transcriptomic landscapes.

Whereas Model A analyses (epitope + IFN-γ versus IFN-γ) identified meaningful disease relevant DEGs only in 65-79*LE- and 65-79*SE-stimulated RAW 264.7 macrophages, Model B analysis (epitope+IFN-γ versus epitope) revealed large numbers of DEGs in cells stimulated by each of the 3 epitopes. (Figs. 2a and 2b). In RAW 264.7 cells, there were 764 unique DEGs in 65-79*LE-treated cells (413 overexpressed and 351 underexpressed), 270 unique DEGs in 65-79*SE-treated cells (156 overexpressed and 114 underexpressed genes), and 449 unique DEGs (244 overexpressed and 205 underexpressed) in 65-79*PE-treated cells (Fig. 2b). Substantial allelic epitope-specificity in gene expression was also found in human THP-1 macrophages (Supplementary Fig.2a).

**Fig. 2:**
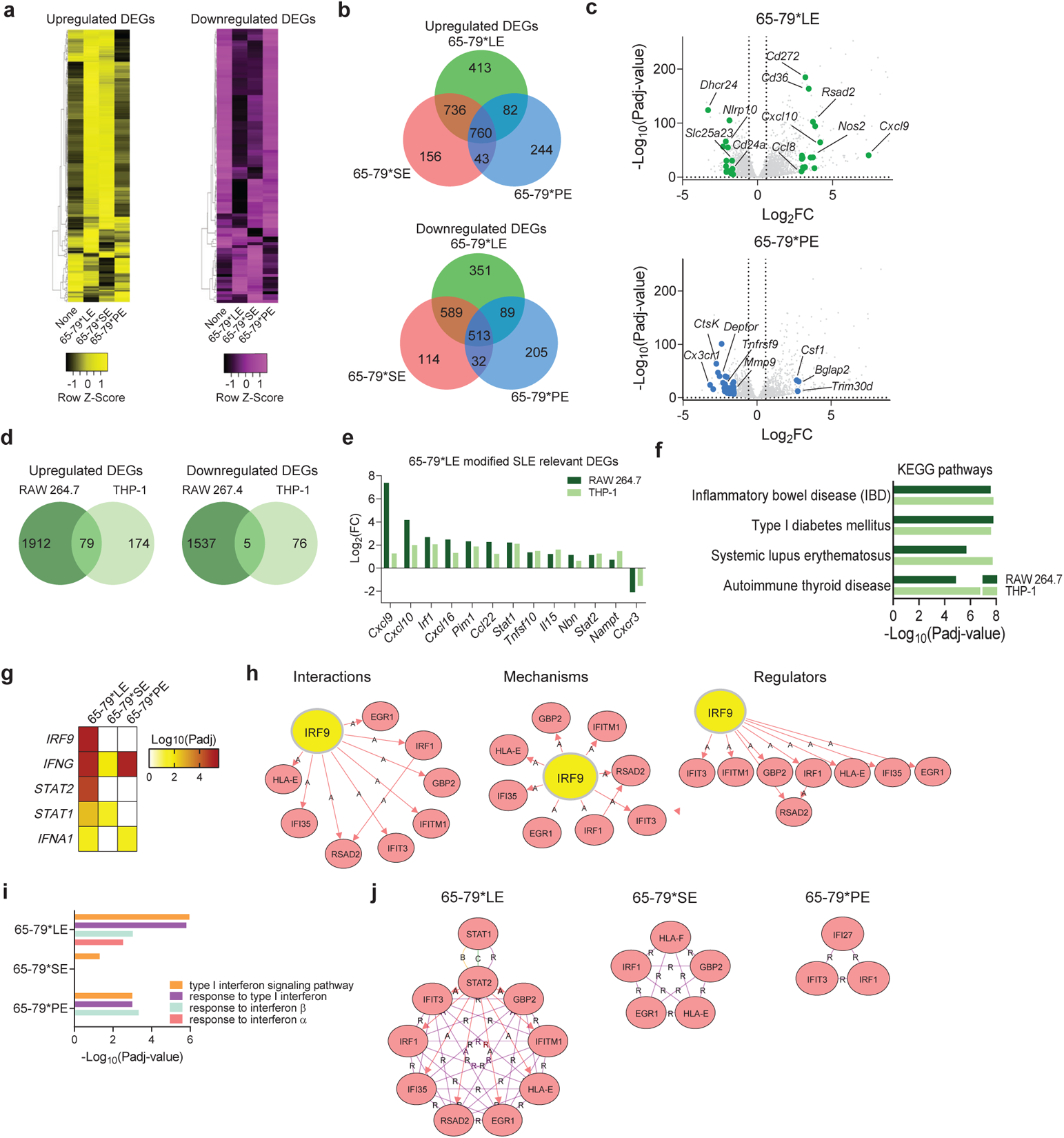
Transcriptional modulation by the LE (Model B: epitope + IFN-γ, versus epitope alone) **a,** Heatmaps showing unsupervised clustering of upregulated (left, yellow) and downregulated (right, purple) DEGs for 65-79*LE, 65-79*SE and 65-79*PE. DEGs were selected based on a > 3-fold change for 65-79*LE. All depicted DEGs had a Padj < 0.05. **b,** Venn diagram comparing DEGs for 65-79*LE, 65-79*SE and 65-79*PE. **c,** Volcano plots comparing the differential transcriptional modulation by 65-79*LE and 65-79*PE. Green dots denote SLE-relevant genes modulated by 65-79*LE; blue dots denote RA-relevant genes modulated by 65-79*PE. **d,** Venn diagrams illustrating the overlap between 65-79*LE-modulated gene expression in RAW 264.7 and THP-1 macrophages. **e,** Representative SLE-relevant genes modulated by 65-79*LE in RAW 264.7 and THP-1 macrophages. **f,** Notable KEGG disease pathway enrichment among 65-79*LE-upregulated DEGs in RAW 264.7 and THP-1 macrophages. **g,** Heatmap of top-ranked URs that are predicted to be activated in 65-79*LE-versus 65-79*SE- and 65-79*PE-modulated transcriptomes in THP-1 macrophages. **h,** Regulatory networks showing the interactions, mechanisms, and regulator modes for IRF9 in THP-1 macrophages. **i,** IFN-I-related GO-BP terms in 65-79*LE-, versus 65-79*SE- or 65-79*PE-activated transcriptomes in THP-1 macrophages. **j,** Comparative hub gene networks for the GO-BP term “type I interferon signaling pathway” in 65-79*LE-, versus 65-79*SE- or 65-79*PE-activated transcriptomes in THP-1 macrophages. Data are based on DEGs with fold change >1.5 and Padj < 0.05. All RAW 264.7 macrophage data are based on 6 biological replicates in 2 independent experiments. THP-1 macrophage data are based on 4 biological replicates. See also Supplementary Fig. 2 and Supplementary Tables 2–4.

In contrast to upregulation of many SLE-relevant genes by 65-79*LE in RAW 264.7 macrophages, 65-79*PE downregulated many pro-inflammatory and pro-SLE genes. Representative genes illustrating the dichotomous modulation of SLE pathogenesis-related gene expression by 65-79*LE versus 65-79*PE are shown in Fig. 2c and Supplementary table 2. In this context, it is notable that *DRB1* alleles that encode a PE motif (70-DERAA-74) have been previously found to significantly reduce the risk of various autoimmune diseases, including SLE^17–19^.

Epitope-activated transcriptional modulation was also evaluated in human macrophages derived from the THP-1 cell line, using Model B, which revealed substantial parallels with the 65-79*LE-activated transcriptomes in mouse RAW 264.7 (Fig. 2d). Among the 79 upregulated DEGs that were shared between the mouse and human macrophage lines, 36 (45.5%) are known to either play roles in SLE pathogenesis, or associate with disease risk (Fig. 2e and Supplementary table 3). All 65-79*LE-modulated DEGs in mouse RAW 264.7 and human THP-1 macrophages are listed in Supplementary Data Files 2 and 3. Additionally, GO-BP analysis identified SLE-relevant terms that were shared between 65-79*LE-treated human and mouse macrophages, including terms pertaining to inflammatory response, response to IFN-γ and type I IFN (IFN-I) signaling pathway (Supplementary Figs. 2b and 2c).

Importantly, KEGG pathway analysis in 65-79*LE-stimulated THP-1 cells (Fig. 2f) ranked several autoimmune diseases, including inflammatory bowel disease (Padj = 1.60×10^−8^), SLE (Padj = 1.83×10^−7^), type I diabetes (Padj = 1.54×10^−7^) and autoimmune thyroiditis (Padj = 7.41×10^−7^) highly. It is noteworthy that allele *DRB1*03:01* has been implicated as a genetic risk factor in all these autoimmune conditions ^9,22–24^.

UR analysis of THP-1 macrophage transcriptomes by iPG (Figs. 2g and 2h) predicted differential involvement of several regulators by the three allelic epitopes. The top predicted URs in 65-79*LE-activated transcriptome included several IFN-I regulators, such as interferon regulating factor 9 (IRF9), as well as two of its interacting proteins, signal transducer and activator of transcription (STAT) 2 and STAT1, previously implicated in SLE pathogenesis ^25–27^. Although STAT1 was predicated to play a regulatory role in both 65-79*LE- and 65-79*SE-activated transcriptomes (Fig. 2g), the two allelic epitopes activated distinct interactions, mechanisms, and regulators (Supplementary Figs. 2d and 2e).

Consistent with the above findings, 65-79*LE-activated THP-1 macrophages showed transcriptional enrichment of multiple IFN-I-associated GO-BP terms (Fig. 2i), and the GO-BP term “Type I interferon signaling pathway” demonstrated an extensive intricate hub gene network in THP-1 macrophages activated by 65-79*LE-compared to 65-79*SE- or 65-79*PE-activated transcriptomes (Fig. 2j).

To identify transcriptional patterns that are shared between Model A and Model B, we performed bioinformatic metanalyses encompassing compiled data sets of RAW 264.7 cells in the two models (Supplementary Fig. 3). A substantial overlap was found between Model A and Model B, with 773 upregulated and 378 downregulated DEGs that were common between the two models in 65-79*LE-stimulated macrophages, and 573 common upregulated and 286 downregulated in 65-79*SE-stimulated macrophages (Supplementary Fig. 3a and 3b). 65-79*PE upregulated only 16 and downregulated no DEGs that were shared by the two models.

GO-BP analysis of upregulated DEGs in 65-79*LE- and 65-79*SE-stimulated macrophages when Model A and Model B were meta-analyzed revealed distinct enrichment patterns between the two epitopes. As shown in Supplementary Figs. 3c and 3d, consistent with the disease relevance of their respective *DRB1*-coding alleles, epitopes 65-79*LE-activated transcriptomes were enriched in terms involving cell division, DNA damage and repair, and nuclear reorganization, whereas transcriptomes in cultures activated by 65-79*SE showed preponderance of GO terms related to proinflammatory and innate immune system signaling, as well as cellular stress response. Additionally, KEGG analysis revealed higher significance among pathways involving cell cycle and SLE in 65-79*LE-stimulated cells, while the higher statistically significant KEGG pathway terms in 65-79*SE-stimulated cells included TNF signaling pathways and several TNF-dependent disease processes, including RA (Supplementary Fig. 3e), consistent with the established disease associations of the two respective coding *DRB1* alleles.

Thus, taken together, RNA-seq analyses identified allele-specific, functionally distinct, and clinically relevant activation of transcriptional landscapes. 65-79*LE activated an SLE-relevant transcriptome, 65-79*SE activated a pro-RA transcriptome, and 65-79*PE activated an autoimmune-protective transcriptome, consistent with the respective disease associations that the three coding *DRB1* alleles demonstrate in humans.

### The LE activates a cascade of lupus-associated cellular aberrations

To determine the functional impact of the three TAHR epitopes we studied their effects in IFN-γ-stimulated mouse RAW 264.7 and human THP-1 macrophages. ER stress and UPR aberrations have been long implicated in SLE pathogenesis ^3,28,29^. As shown in Fig. 3a, RNA-seq analyses identified many ER stress and proteasome degradation genes that were differentially regulated by 65-79*LE. We therefore determined the functional impact of 65-79*LE on these cellular processes. In the presence of IFN-γ, THP-1 cells activated by 65-79*LE, but not by 65-79*SE or 65-79*PE, showed increased expression of the ER stress marker phosphorylated inositol-requiring enzyme 1 α (pIRE1α), C/EBP homologous protein (CHOP) and glucose regulated protein 78 (GRP78) (Fig. 3b), as well as the proteasomal degradation marker p62 and overabundant poly-ubiquitinated proteins (Figs. 3c, 3d and 3e). Evidence of allele-specific 65-79*LE-triggered ER stress and UPR was found in RAW 264.7 macrophages as well (Supplementary Fig. 4).

**Fig. 3:**
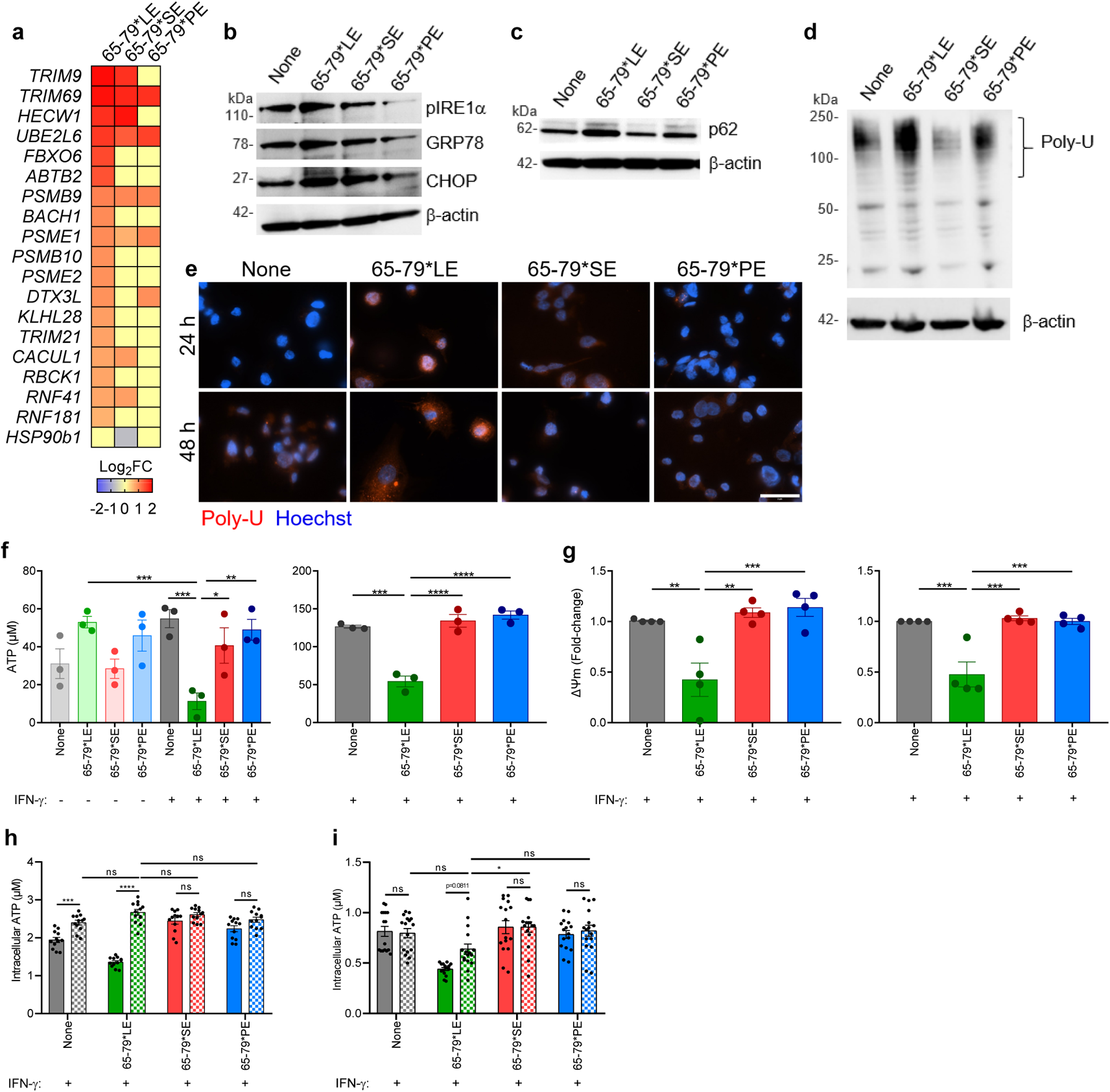
Activation of ER stress, UPR and mitochondrial dysfunction by the LE in human and mouse macrophages. **a,** Heatmap of endoplasmic reticulum-associated degradation pathway-related genes (Model B) modulated by 65-79*LE, versus 65-79*SE and 65-79*PE in IFN-γ-treated THP-1 macrophages. **b** – **d**, Representative immunoblots of ER stress markers (pIRE1-α, GRP78, CHOP) **b,** proteasomal degradation marker (p62) **c,** and poly-ubiquitinated proteins **d,** in THP-1 macrophages in the presence of 65-79*LE, 65-79*SE, or 65-79*PE. **e,** Representative immunocytochemistry images of poly-ubiquitinated protein accumulation (red). Scale bar = 2 μm. **f,** Intracellular ATP levels reduction in allelic epitope-treated human THP-1 (left) and mouse RAW 264.7 (right) macrophages (n=3). **g,** Mitochondrial membrane potential in allelic epitope-treated THP-1 (left) and RAW 264.7 (right) macrophages, as determined by tetramethylrhodamine ethyl ester (TMRE), red (n=4). **h**, Baricitinib (10 μg/mL) shows allele-nonspecific inhibitory effect on LE-activated reduction of intracellular ATP in RAW 264.7 macrophages, (n=3). **i**, Neutralization of human IFN-alpha receptor with IFNAR2 Monoclonal Antibody (10 μg/mL) shows a trend for recovery of LE-activated intracellular ATP levels, in THP-1-derived macrophages, (n=3). In **h,i**, solid-color and dotted-color bars represent, respectively, absence or presence of inhibitors. Data represent mean ± SEM; One-way (**f,g),** or two-way (**h,i**) ANOVA.*p < 0.05; **p < 0.01; ***p < 0.001, ****p < 0.0001. See also Supplementary Fig. 4.

The ER and mitochondria are functionally entangled ^30,31^. We therefore sought to characterize indicators of mitochondrial function, including ATP levels, mitochondrial membrane potential (ΔΨm) and mitochondrial reactive oxygen species (ROS) levels. In the presence of IFN-γ, macrophages stimulated with 65-79*LE, but not with 65-79*SE, 65-79*PE, or with 2 other 15mer peptides corresponding to the TAHR coded by other control alleles, *DRB1*04:03* and *DRB1*15:01,* demonstrated significantly, and dose-dependently, decreased cellular ATP levels (Fig. 3f and Supplementary Fig.1). Corroborating evidence for 65-79*LE-activated mitochondrial dysfunction was the finding of an allele-specific effect of this epitope on mitochondrial membrane potential. In the presence of IFN-γ, 65-79*LE, but not 65-79*SE or 65-79*PE, significantly suppressed mitochondrial membrane potential and increased mitochondrial ROS both in human THP-1 and mouse RAW 264.7 macrophages (Figs. 3f and 3g and Supplementary Fig. 4). Reduction of intracellular ATP levels by 65-79*LE could be completely prevented in THP-1 macrophages by Baricitinib (Fig. 3h), a Janus Kinase inhibitor known to block IFN-I-activated pathways ^32^. Additionally, a clear trend toward ATP normalization was found in cultures treated with a monoclonal antibody against IFN-I receptor, IFNAR2, as well (Fig. 3i). Thus, IFN-I appears to play a role in 65-79*LE-activated cell aberrations, consistent with the IFN-I-associated GO-BP terms enrichment effects by this epitope (Figs. 2i and 2j), and the known role of IFN-I in SLE.

Since mitochondrial dysfunction is a key factor involved in cell death aberrations ^35,36^, which have long been implicated in SLE pathogenesis ^2,37^, we next sought to determine if 65-79*LE-activated mitochondrial dysfunction leads to cell death. In the presence of IFN-γ, 65-79*LE-treated human THP-1 macrophages, but not macrophages treated with 65-79*SE or 65-79*PE, showed accelerated cell death (Fig. 4a), and reduced metabolic activity (Fig. 4b). Increased ROS levels could be seen in THP-1 macrophages stimulated with either 65-79*LE or 65-79*SE, but not with 65-79*PE (Fig. 4c). Consistent with an RNA-seq-based DNA damage gene expression heatmap (Fig. 4d), in the presence of IFN-γ, 65-79*LE facilitated DNA damage in THP-1 macrophages in an allele-specific fashion (Fig. 4e).

**Fig. 4:**
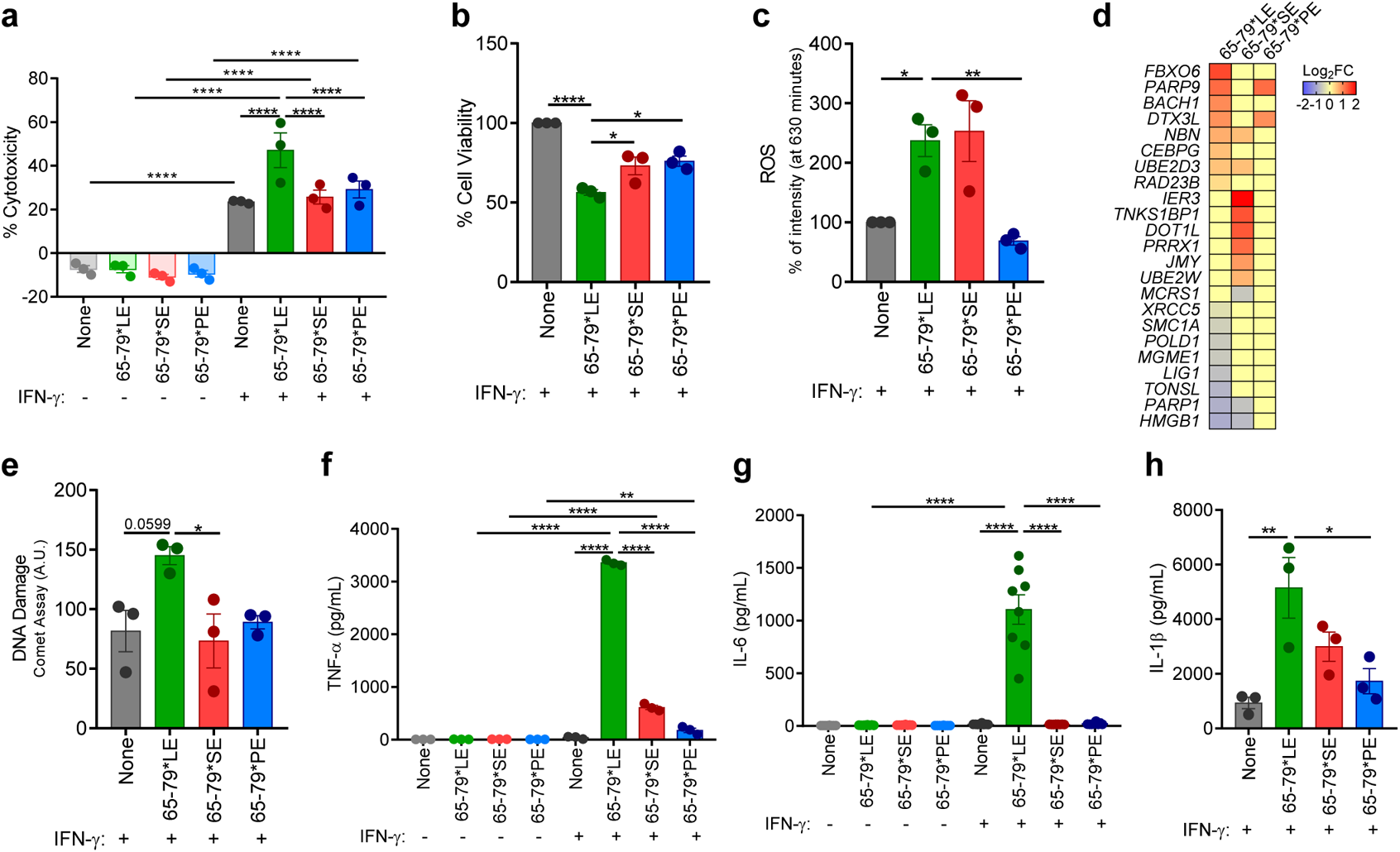
Allele-specific activation of cell death, pro-inflammatory cytokine production and DNA damage by the LE. **a-c**, Cell death **a,** viability **b,** and ROS levels **c,** in THP-1 cells exposed to different allelic epitopes (n=3). **d,** Heatmap of DNA damage-related genes (Model B) in THP-1 cells exposed to different allelic epitopes (n=4). **e,** DNA damage, quantified by the comet assay, in THP-1 cells exposed to different allelic epitopes (n=3). **f**-**h**, Supernatant levels of the pro-inflammatory cytokines TNF-α **f,** IL-6 **g,** and IL-1β **h,** in THP-1 macrophages exposed to different allelic epitopes (n=3). Data represent mean ± SEM. One-way (**b**, **c**, **e**, **h**), or two-way (**a**, **f**, **g**) ANOVA. *p < 0.05, **p < 0.01, ***p < 0.001, ****p < 0.0001. See also Supplementary Fig. 5.

Importantly, 65-79*LE-facilitated cell death was associated with a robust release of pro-inflammatory cytokines (Fig. 4f-4h). Similar 65-79*LE-activated release of proinflammatory mediators was observed in mouse RAW 264.7 macrophages, as well as primary, bone marrow-derived macrophages (BMDMs) isolated from 3 genetically distinct wild type (WT) mouse strains (Supplementary Fig. 5). Cell death induced by 65-79*LE did not indicate involvement of caspase 3 activation and was resistant to inhibition by a pan-caspase inhibitor, ZVAD-FMK (Fig. 5 and Supplementary Fig.6). Additionally, there was no significant evidence for autophagy, and rapamycin, an autophagy inducer had no effect on 65-79*LE-activated cell death in THP-1 cells, or allele-nonspecific effects in RAW 264.7 (Fig. 5 and Supplementary Fig. 6).

**Fig. 5:**
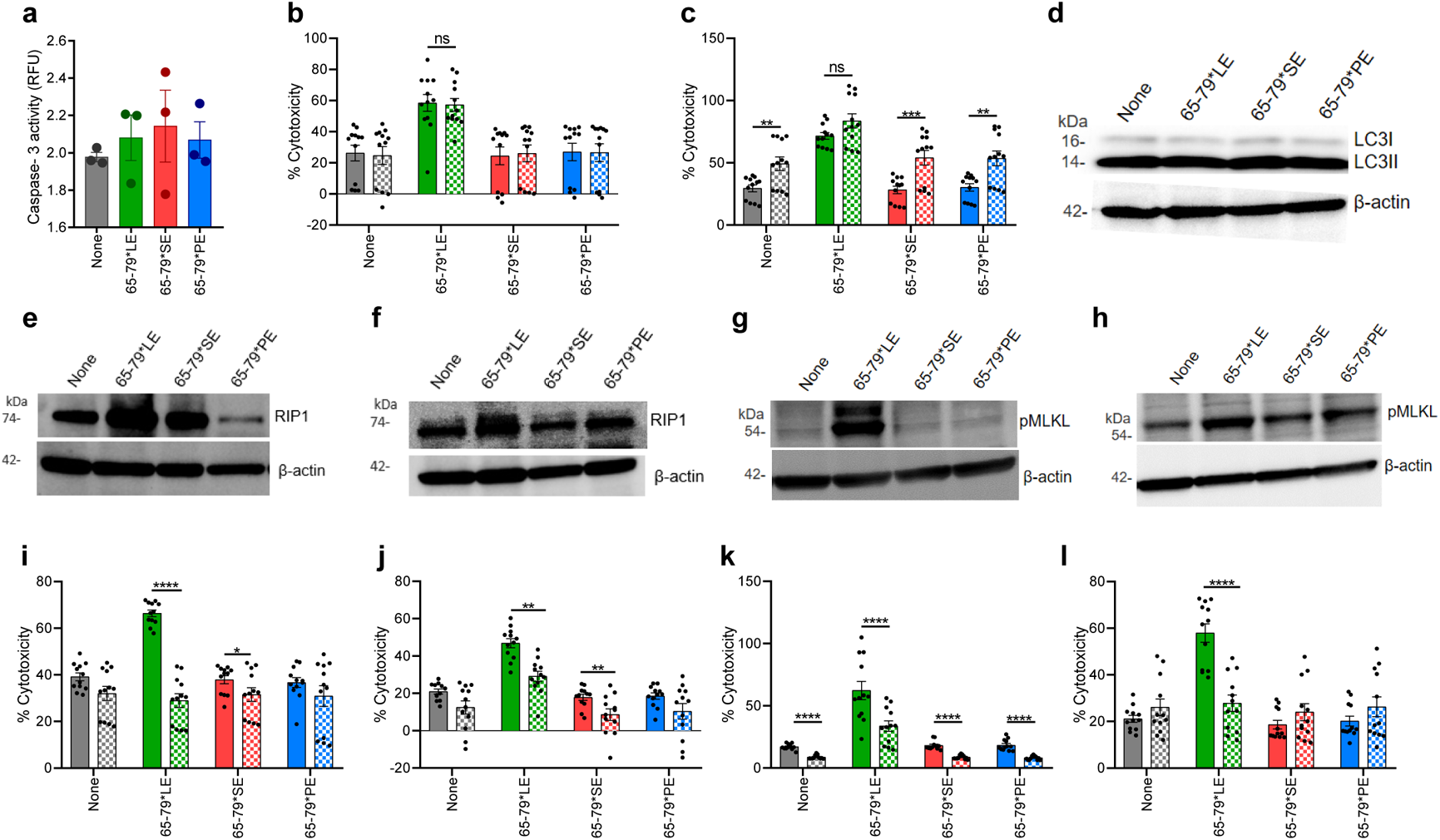
The LE triggers necroptotic cell death in an allele-specific fashion. **a,** Caspase-3 activity in THP-1 macrophages (n=3). **b,** Cell death in THP-1 macrophages in the absence (solid bars) or presence (dotted bars) of pan-caspase inhibitor ZVAD-FMK (10 µM) (n=3). **c,** Cell death in THP-1 macrophages in the absence (solid bars) or presence (dotted bars) of rapamycin (50 nM) (n=3). **d,** Immunoblot of LC3 in RAW 264.7 macrophages. **e,** Immunoblot of Receptor-Interacting Protein 1 (RIP) in RAW 264.7 macrophages. **f,** Immunoblot of RIP in THP-1 macrophages. **g,** Immunoblot of pMLKL in RAW 264.7 macrophages. **h,** Immunoblot of pMLKL in THP-1 macrophages. **i**, Cell death of RAW 264.7 macrophages in the presence (dotted bars) or absence (solid bars) of the RIP1 inhibitor Necrostatin-1 (50 μM) (n=3). **j**, Cell death of THP-1 macrophages in the presence (dotted bars) or absence (solid bars) of the RIP1 inhibitor Necrostatin-1 (50 μM) (n=3). **k**, Cell death of RAW 264.7 macrophages in the presence (dotted bars) or absence (solid bars) of the MLKL inhibitor Necrosulfonamide (10 μM) (n=3). **l**, Cell death of THP-1 macrophages in the presence (dotted bars) or absence (solid bars) of the MLKL Inhibitor Necrosulfonamide (10 μM) (n=3). Blots (**d**-**h**) are representative of 3 independent experiments each. Data represent mean ± SEM; One-way **a,** or two-way (**b**, **c**, **i**-**l**) ANOVA. *p < 0.05, **p < 0.01, ***p < 0.001, ****p < 0.0001. See also Supplementary Fig. 6.

We proceeded to determine whether 65-79*LE-activated cell death involves necroptosis, an inflammatory, caspase-independent cell death ^38,39^, which has been shown to feature in SLE ^40^. Indeed, 65-79*LE-treated macrophages showed increased expression of the necroptosis markers receptor-interacting serine/threonine-protein 1 (RIP1) and phosphorylated mixed lineage kinase domain like pseudo kinase (pMLKL) (Figs. 5e-5h). Moreover, the RIP1 inhibitor necrostatin1 (Nec-1) and the MLKL inhibitor necrosulfonamide (NSA) blocked potently and allele-specifically 65-79*LE-activated cell death (Figs. 5i-5l).

Crosstalk between mitochondria and ER has been previously documented, with ample evidence indicating that mitochondrial dysfunction is often secondary to ER stress ^41–43^. To determine whether the mitochondrial dysfunction observed in 65-79*LE-treated macrophages is secondary to ER stress, we used sodium 4-phenylbutyrate (4-PBA), a chemical chaperone that inhibits protein misfolding, thereby inhibiting ER stress ^44^. Relevant to the present study, 4-PBA has been shown to prevent SLE manifestations in experimental lupus models ^45,46^. As expected, 4-PBA inhibited 65-79*LE-activated ER stress and proteasomal degradation (Fig. 6a). Further, prevented reduction of intracellular ATP levels (Fig. 6b), reversed mitochondrial membrane potential loss (Fig. 6c), and normalized mitochondrial ROS levels (Fig. 6d), in an allele-specific fashion. Moreover, 4-PBA inhibited 65-79*LE-activated necroptosis signaling as evidenced by decreased expression of the necroptotic protein markers RIP1 and pMLKL, as well as cell death in mouse and human macrophages (Figs. 6e-6g).

**Fig. 6:**
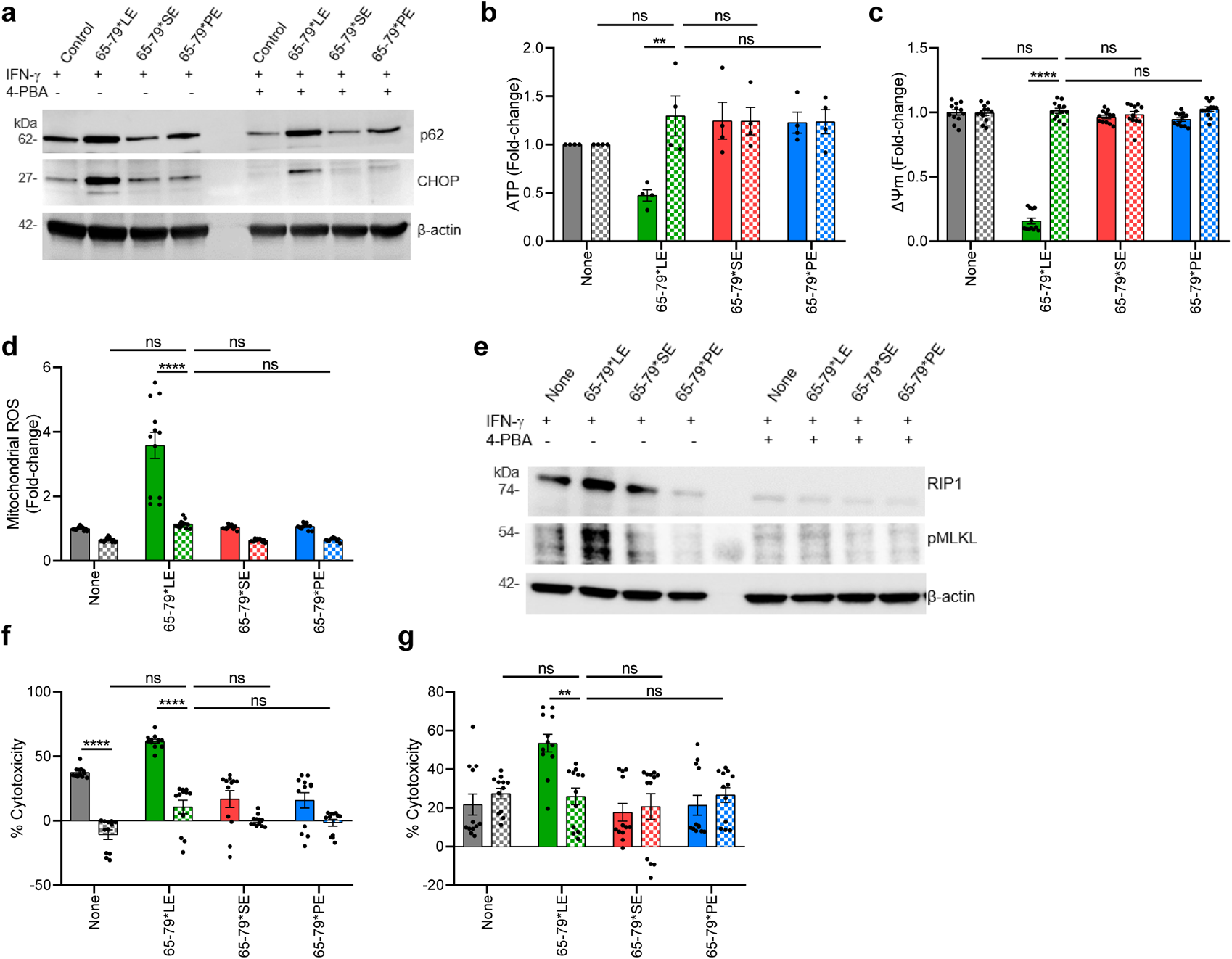
LE-activated mitochondrial dysfunction is ER stress-dependent. **a,** Immunoblots of CHOP and p62 in allelic epitope-treated RAW 264.7 macrophages in the presence (dotted bars) or absence (solid bars) of the ER stress inhibitory chemical 4-PBA (5 mM). **b**,**d**, Intracellular ATP levels **b,** mitochondrial membrane potential **c,** and mitochondrial ROS levels **d,** in RAW 264.7 macrophages in the presence (dotted bars) or absence (solid bars) of 4-PBA (n=3). **e,** Immunoblots of RIP1 and pMLKL in allelic epitope-treated RAW 264.7 macrophages in the presence (dotted bars) or absence (solid bars) of 4-PBA. **f**,**g**, Cell death in allelic epitope-treated RAW 264.7 **f,** and THP-1 **g,** macrophages in the presence (dotted bars) or absence (solid bars) of 4-PBA (n=3). Blots (**a**, **e**) are each a representative of 3 independent experiments. Data represent mean ± SEM. Two-way ANOVA, **p < 0.01, ****p < 0.0001. See also Supplementary Fig. 6.

To better map the specific pathway involved, we tested three selective inhibitors that target the transcription factor 6α (ATF6α)-, protein kinase R-like endoplasmic reticulum kinase (PERK)- or inositol-requiring enzyme-1 (IRE1)-mediated pathways (Supplementary Fig. 6h). The findings suggest that inhibition of the ATF6α-, but not PERK- or IRE1-mediated pathways significantly increased cellular ATP levels, suggesting that that LE-activated UPR pathway is mediated by ATF6α (Supplementary Fig. 6i-6k).

### Disease-promoting effects of physiologically folded LE-expressing HLA-DR molecules in transgenic mice

To determine the functional effects of the LE in its physiologically folded conformation, we performed *ex vivo* and *in vivo* experiments in transgenic mice that express on their cell surfaces heterodimeric HLA-DR molecules consisting of a monomorphic DRα chain and a polymorphic DRβ chains coded by one of the three alleles *DRB1*03:01* (LE); *DRB1*04:01* (SE), or *DRB1*04:02* (PE). These transgenic mouse lines are referred to as LE-Tg, SE-Tg and PE-Tg, respectively. Their cell surface-expressed HLA-DR molecules differ from each other by a few amino acid residues in the 65-79 region of the DRβ chain (Supplementary Fig. 1b).

Genomic DNA analyses of the three Tg mouse lines using the MiniMUGA SNP array showed that the three Tg mouse lines include identical background strain compositions, with minor difference in the calculated percentage of genomic contribution by C57BL/6J (B6J), C57BL/10J (B10J) and SWR mouse strains (detailed in the Methods section). We therefore used these 3 WT mouse strains as genetic background controls in key experiments.

Consistent with the findings in mouse and human macrophage cell lines discussed above, when exposed to IFN-γ, primary bone marrow-derived macrophages (BMDMs) from LE-Tg showed evidence of ER stress and UPR activation (Figs. 7a and 7b), and reduction of intracellular ATP levels (Fig. 7c). Such effects could not be seen in LE-Tg BMDM in the absence of IFN-γ, nor could they be seen in the presence or absence of IFN-γ, in the control, SE-Tg or PE-Tg BMDMs, or in the 3 genetic background control WT mouse strains, B6J, B10J and SWR (Supplementary Fig. 7). Like the effect of 65-79*LE on mouse and human macrophage cell lines (Fig. 4 and Supplementary Fig. 5), in the presence of IFN-γ, BMDMs from LE-Tg showed increased proinflammatory milieu, as evidenced by increased production of TNF-α and nitrite (Figs. 7d and 7e). Such proinflammatory effects could not be seen in LE-Tg BMDM in the absence of IFN-γ,

**Fig. 7:**
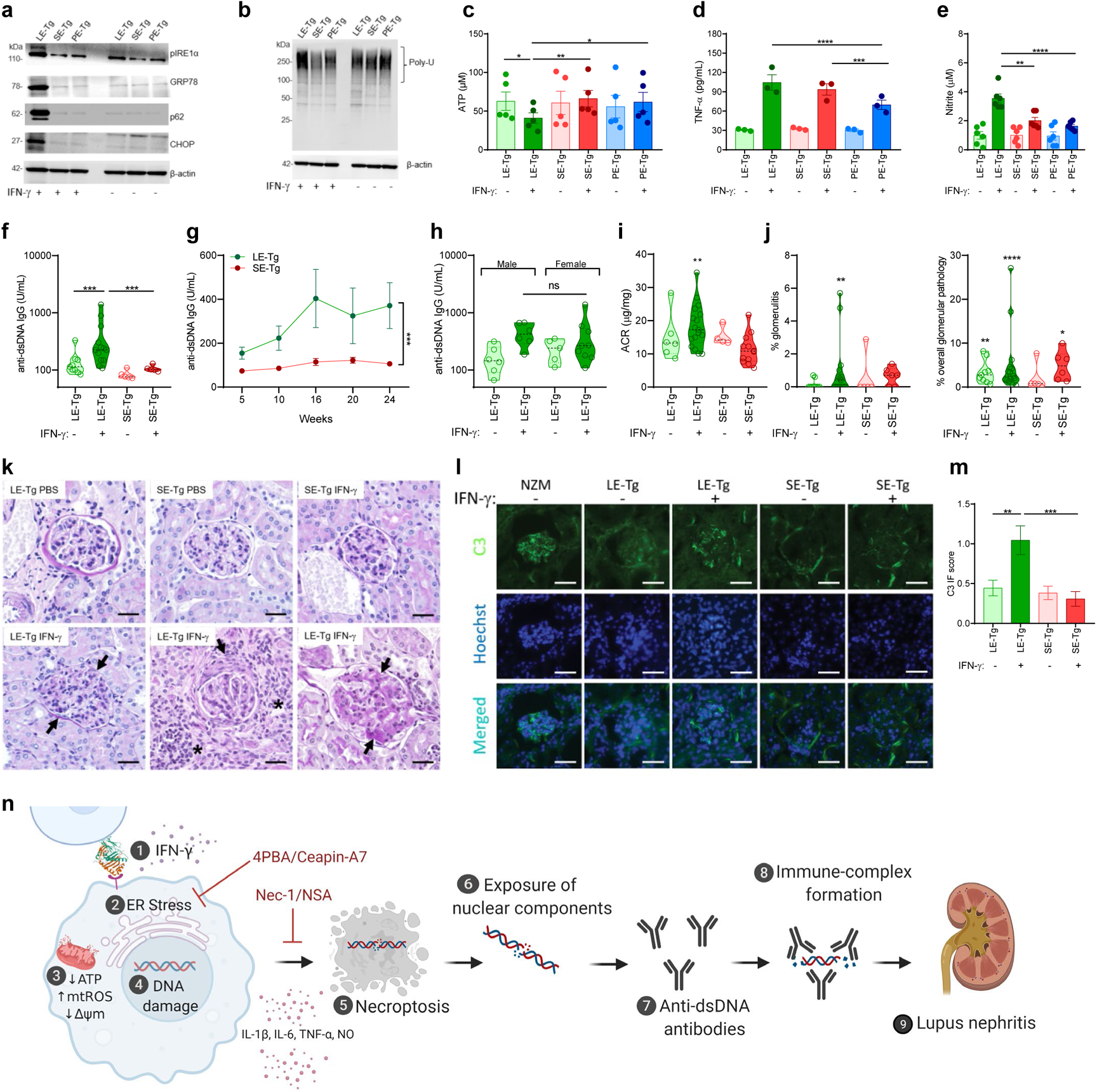
*Ex vivo* and *in vivo* effects of physiologically folded LE-expressing HLA-DR molecules. **a**, **b**, Immunoblots of ER stress (pIRE1-α, CHOP, GRP78) and proteasomal degradation (p62) markers **a,** and poly-ubiquitinated proteins **b,** in BMDMs derived from transgenic mice expressing physiologically folded HLA-DRβ molecules coded by *DRB1*03:01* (LE-Tg), *DRB1*04:01* (SE-Tg), *or DRB1*04:02* (PE-Tg) and cultured *ex vivo* in the presence or absence of IFN-γ (5ng/mL). **c**-**e**, Intracellular ATP **c,** TNF-α **d,** and nitrite **e,** supernatant levels in BMDMs cultured as in (**a**,**b**). (**f**-**l**) Groups of LE-Tg and SE-Tg were administered 10 μg recombinant mouse IFN-γ or PBS, twice weekly intraperitoneally over a 24-week period. **f,** Serum anti-dsDNA IgG levels measured at week 24 in LE-Tg (treated with PBS, n=9, or IFN-γ, n=13), and SE-Tg (treated with PBS, n=5, or IFN-γ, n=5) mice. **g,** Time-course curves of serum anti-dsDNA IgG levels in IFN-γ-treated LE-Tg (n=9-13) and SE-Tg (n=5-6). **h,** Comparison of anti-dsDNA IgG levels in male (n=6) and female (n=9) LE-Tg mice treated for 16 weeks with IFN-γ. **i,** Albumin-to-creatinine ratio (ACR) in LE-Tg (PBS, n=6, IFN-γ, n=17) and SE-Tg (PBS, n=4, IFN-γ, n=12) mice. **j**, Percent of renal glomeruli showing microscopic evidence of glomerulitis (left) or overall glomerular pathology (right) in LE-Tg (PBS, n=9, IFN-γ, n=15) and SE-Tg (PBS, n=5, IFN-γ, n=6) mice. **k**, Representative images of PAS-stained kidney tissues from PBS- or IFN-γ-treated LE-Tg and SE-Tg mice. Arrows point at illustrative examples of endocapillary hypercellularity (lower left), fibrous crescent (lower middle) and immune deposits (lower right). Asterisks identify interstitial inflammation areas. Scale bar = 20 µm. **l**, Representative images of glomerular C3 deposition patterns in PBS- or IFN-γ-treated LE-Tg and SE-Tg mice. Scale bar = 50 µm. **m**, C3 immunofluorescence scoring. **n**, A proposed model of the LE-activated pathogenetic cascade. In the presence of IFN-γ [1], the LE – possibly through interaction with a yet unknown cell surface receptor - triggers ER stress [2], which leads to mitochondrial dysfunction [3], DNA damage [4], and necroptosis [5]. Concomitantly released pro-inflammatory cytokines enhance the immunogenicity of nuclear antigens [6], such as dsDNA, with resultant increased levels of anti-dsDNA antibodies [7] and formation of immune complexes [8], which are deposited in the kidneys [9]. Immunoblots (**a** and **b**) are representatives of 3 independent experiments each. Data represent mean ± SEM. One-way ANOVA (**c** - **e**, **g**, **h**, **l**), or Wilcoxon sign rank test (**f**, **i**). *p < 0.05, **p < 0.01, ***p < 0.001, ****p < 0.0001.

To ascertain that BMDMs from the three genetic background control strains are capable of responding to an exogenously added LE epitope, we measured their proinflammatory response to the allelic peptides in the presence or absence of IFN-γ. Interestingly, like mouse RAW 264.7 and human THP1 macrophage cell lines, in the presence of IFN-γ, BMDM of all the three control mouse strains (B6J, B10J and SWR) showed increased TNF-α and nitrite production in response to exogenously-added LE peptide 65-79*LE, but not to 65-79*SE or 65-79*PE (Supplementary Figs. 5f and 5g). These findings demonstrate that LE-triggered proinflammatory effects are strain- and species-independent.

Given our findings of an LE-activated IFN-I transcriptome (Fig. 2 and Supplementary Fig. 2), and the known pathogenic role of IFN-I in both human SLE and mouse models of the disease ^47^, we quantified by qPCR the expression levels of salient IFN-I genes in HLA-DR humanized mice expressing on their surface LE-, or SE-positive HLA-DR molecules (LE-Tg and SE-Tg mice, respectively). As supplementary Fig. 8 demonstrates, in the presence of IFN-γ, BMDMs from LE-Tg mice, but not from control, SE-Tg mice, showed increased gene expression of several known SLE-associated IFN-I genes, such as *Ifi44, Ifi44l,* and *Irf7*, along with markedly curtailed expression of *Dnase1l3*, a gene known to protect against SLE in both humans and mice^48–50^.

IFN-γ has been long found to contribute to SLE disease development in both humans and mice ^15,16^. Given the observed obligatory co-factor role of IFN-γ *in vitro* at the transcriptome and cell function levels as noted above, we proceeded to determining its *in vivo* effect in Tg mice. Intraperitoneal administration of IFN-γ resulted in increased serum levels of anti-dsDNA antibodies in LE-Tg, but not in SE-Tg mice (Figs. 7f and 7g), without a significant difference between male and female mice (Fig. 7h). IFN-γ-treated LE-Tg mice developed glomerulonephritis, as evidenced by proteinuria (Fig. 7i) and histopathology (Figs. 7j and 7k). Specifically, examination of kidney sections of LE-Tg mice treated for 24 weeks with IFN-γ showed increased glomerulitis with mesangial and/or endocapillary hypercellularity, crescent formation and immune deposits, as compared to LE-Tg mice treated with PBS, or SE-Tg mice treated with PBS or IFN-γ (Fig. 7k). In addition, kidney sections of LE-Tg mice treated with IFN-γ showed increased glomerular C3 deposits compared to LE-Tg mice treated with PBS, or SE-Tg mice treated with either PBS or IFN-γ (Figs. 7l and 7m).

Thus, taken together, the findings of this study demonstrate that in the presence of IFN-γ, the LE activates lupus signature transcriptomes and a cascade of lupus-associated cellular events *in vitro* in an allele-specific manner. When exposed to IFN-γ *in vivo*, naïve transgenic mice carrying the LE-coding SLE risk allele *DRB1*0301* develop serological and histological findings analogous to human SLE. A proposed model is illustrated graphically in Fig. 7n.

## DISCUSSION

It has been long proposed that the mechanistic basis of HLA-disease association in lupus involves presentation of putative self or foreign antigens that lead to an adaptive immune system-mediated target organ damage. The new insight offered by this study is that an entire cascade of SLE-associated events, including transcriptional, cellular, serological, and clinical manifestations, can be triggered, independent of antigen presentation, by a short epitope coded by allele *DRB1*03:01*, known as a major risk factor in human SLE.

Studying mouse and human macrophage cell lines stimulated with short synthetic peptides corresponding to the TAHR coded by three *DRB1* alleles, as well as BMDMs derived from transgenic mouse lines that express physiologically folded distinct allelic HLA-DR molecules, we demonstrated allele-specific activation of a cascade of pathogenic events that includes transcriptional, cellular and disease phenomes that closely resemble human SLE and its experimental mouse models. Given that synthetic peptides are incapable of presenting antigens, and the fact that macrophage cell cultures are devoid of T cells, these findings support a disease-relevant, *DRB1* allele-specific, antigen presentation-independent mechanism.

Consistent with its known effects in human SLE and experimental models of the disease ^15,16^, IFN-γ was identified here as an obligatory co-factor for LE-activated transcriptional and cell biology effects. In this context, it is of interest that while both Model A (epitope + IFN-γ versus IFN-γ alone) and Model B (epitope + IFN-γ versus epitope alone) of RNA-seq data analysis uncovered bioinformatics evidence of a lupus signature transcriptome activation by the LE, Model B was found to be more robust in both mouse and human macrophages. One possible explanation might be that Model B more closely mimics the physiologic conditions *in vivo*, where the expression levels of *DRB1*-coded HLA-DR molecules is relatively constant, while IFN-γ levels could vary periodically, secondary to fluctuating environmental conditions. It is plausible that such changing circumstances could allow the LE to realize its SLE-triggering potential. Further research into this question, and more detailed mapping of the LE pathway and its crosstalk with IFN-γ-activated pathways, will help to dissect the respective roles of LE and IFN-γ and the mechanism of their interaction in SLE.

The LE overcame strain and species barriers, as the effects of this human gene product could be found in mouse (RAW 264.7 and BMDMs of diverse mouse strains) as well as human (THP-1) cells. Additionally, LE effects were found both in cell lines (RAW 264.7 and THP-1) and primary cells (BMDMs). This versatility attests to the fundamental nature of the mechanisms involved in LE-activated events. It is noteworthy that the MHC Cusp theory discussed below is partly based on the hypothesis that class II β chain TAHRs may have preserved fundamental, non-antigen presenting functions through evolution, consistent with the principles of trans-species polymorphism ^11,51,52^.

Different from the strong female preponderance in human SLE, here, LE effects on anti-dsDNA antibody levels showed no difference between male and female mice. The reason for this apparent discrepancy is unknown and requires further study; however, it is worth mentioning that although a higher incidence and severity of SLE disease is well documented in female mice in some experimental models such as NZBW ^53^, sex bias is less prominent in the MRL/lpr model ^54^. Notably, a large genome-wide association study involving genotype data from over 27,000 individuals identified no gender based differential SLE risk for any HLA allele ^55^, suggesting that gender bias in humans involves mechanisms influenced by non-HLA risk factors.

The penetrance of nephritis in IFN-γ-treated LE-Tg mice was low, but statistically significant. It is worth noting that anti-dsDNA antibody formation, proteinuria histopathology and immunofluorescence findings, specifically in LE-Tg mice, were all consistent with LN development. We submit that LE-Tg + IFN-γ may not be a robust model of murine SLE, and there are several popular models with higher penetrance and disease severity. However, our LN data are not presented to propose a new robust lupus model, but rather to substantiate the disease relevance of the *in vitro* data. Susceptibility and disease severity in SLE are multifactorial. It might be helpful to explore in future studies whether the LE can act as a disease facilitator in established models of SLE.

This study has focused on *DRB1*03:01*, the best characterized and the single most significant (OR = 1.87; *p* = 1.17×10^−58^) ^9^ SLE-associated HLA allele. However, it should be mentioned that SLE has been reported to associate with other HLA genes as well. While some of these associations are attributable to the strong linkage disequilibrium in the MHC region, *DRB1*15:01* has been found to independently associate with the disease as well (OR=1.33; *p* = 1.92×10^−12^) ^9^. The mechanistic basis of that association is unknown and is beyond the scope of this first study. It is worth noting, however, that a recent association study of SLE using Immunochip genotyping data ^55^ identified differential risk associations at a single amino acid residue level between the *DRB1*15* allele, in which risk-associated residues were found primarily inside the peptide-binding pocket, and allele *DRB1*03:01*, in which risk-associated residues were found primarily at the region 70-77 of the DRβ chain, overlapping with the cusp region (amino acid residues 65-79).

The distinct topologies of SLE-critical amino acid residues coded by the two alleles, along with our findings that the 65-79*LE, but not an equivalent peptide corresponding to *DRB1*15:01*, suppressed ATP cellular levels (Supplementary Fig.1d), together suggest that the functional effects on disease risk by *DRB1*03:01* and *DRB1*15:01* are dissimilar. While allele *DRB1*03:01* encodes a disease-driving LE, as demonstrated here, allele *DRB1*15:01* may contribute to disease risk through antigen presentation. The apparent disparities between the respective functional effects of the two alleles on disease risk might also provide a mechanistic basis for an imputation-based conclusion that *DRB1*03:01/DRB1*15* heterozygosity confers a greater risk for SLE than either *DRB1*03:01/DRB1*03:01* or *DRB1*15/DRB1*15* homozygosity^55^.

Different from the empirical data-based model presented here, the above-mentioned gene association study suggested that SLE disease risk conferred by allele *DRB1*03:01* is attributable to a single amino acid residue located at the peptide binding groove: Tyr26 in the risk allele *DRB1*03:01* versus Phe26 in the non-risk allele, *DRB1*03:02*, indirectly implicating antigen presentation. That conclusion was offered because both alleles share an identical amino acid sequence in the TAHR but differ by a single amino acid residue in position 26, located at the DR3 peptide groove ^55^.

That hypothesis is seemingly in conflict with our findings. However, despite continuous refining efforts, the accuracy of single nucleotide polymorphism-based imputation of HLA genotyping, especially in respect to the *DRB1* locus ^56,57^, remains uncertain. Nonetheless, irrespective of the reliability of computational HLA fine genotyping, the theory examined here and the antigen presentation hypothesis as mechanistic bases for HLA-disease association are mutually non-exclusive, as previously discussed ^10,11,20,58^. Indeed, targeted sequencing efforts to examine the genetic association within the HLA class II region in lupus identified a regulatory genetic variant located between *HLA-DRB1* and *HLA-DQA1* that could explain the majority of the genetic risk in this locus ^59^. This effect is tagged by the SNP rs9271593 within an intergenic regulatory locus known as *XL9* separating the *HLA-DRB1* and *HLA-DQA1* genes. The lupus-associated variant in this SNP tags the lupus-associated classical alleles *DRB1*03:01* and *DRB1*15:01*, and is associated with increased expression of HLA-DRB1, HLA-DQA1, and HLA-DQB1 in monocyte-derived dendritic cells ^59^. These findings support an antigen presentation independent effect of the HLA class II genetic effect in lupus, consistent with our findings.

It should be noted that *DRB1*03:01* is associated with many other autoimmune diseases besides lupus. We submit that as demonstrated here, the LE activates several fundamental cell biology aberrations (e.g., UPR, mitochondrial perturbations), which are implicated in many diseases besides SLE. Additionally, SLE risk, like many other autoimmune conditions, depends on multiple genes, as well as environmental factors and disease phenomes are likely determined by intricate gene-gene and gene-environment interactions. The fact that KEGG pathway transcriptome analyses of LE-activated macrophages identified several other diseases that are known to associate with *DRB1*03:01* in both human and mouse macrophages (Fig. 2f) lend further support to the mechanism proposed here.

This study demonstrates an allele-specific, antigen presentation-independent cascade of events that culminate in development of serological and tissue damage akin to human SLE. The trigger of this cascade of events involves a short epitope coded by *HLA-DRB1*03:01*, the single most significant genetic risk factor for SLE in diverse populations. Our model (Fig. 7n) proposes that a combination of necroptotic cell death with increased abundance of free DNA fragments plus a proinflammatory milieu, together stimulate anti-dsDNA antibody formation. The model further proposes that those antibodies can form immune complexes with DNA fragments and precipitate in glomeruli, and cause LN. Importantly, the curtailed IFN-γ-activated transcription of *Dnase1l3*, - a gene that codes for an extracellular DNA degradation enzyme, whose deficiency is strongly implicated in the pathogenesis of human SLE ^50^ and murine disease models ^49^ - in BMDM of LE-Tg (Supplementary Fig. 8) suggests a possible facilitating mechanism of anti-dsDNA antibody formation by increasing antigenic (DNA) exposure. This scenario deserves further research.

The findings of this study might open the door to better understanding of the precise molecular and genetic mechanisms involved in HLA-disease association. Exploration of such mechanisms in the context of human diseases could potentially identify new preventive, diagnostic, and therapeutic solutions.

## METHODS

### Mice

Transgenic (Tg) mice were generated as previously described ^24,60,61^ and donated by Dr. Chella David and Dr. Veena Taneja from the Mayo Clinic. These Tg mouse lines, expressing cell surface heterodimeric HLA-DR molecules made of a monomorphic DRα chain and a polymorphic DRβ chain coded by the human *HLA-DR1*0301*, *HLA-DR4*04:01* or *HLA-DR4*04:02* alleles, are referred to as LE-Tg, SE-Tg and PE-Tg, respectively.

Transgenic lines LE-Tg, SE-Tg and PE-Tg were developed using classical pronuclear microinjection. For line LE-Tg, the transgene was injected into (C57BL/6 x DBA/2)F1 embryos ^62^. Lines SE-Tg and PE-Tg were created injecting (C57BL/10 x SWR)F1 embryos ^63,64^. Genomic DNA from four representative mice from each line was analyzed using the MiniMUGA SNP array, containing ~9,700 SNPs ^65^. The three lines showed to be highly inbred (98.5% to 99.5% homozygous genotypes) and mice from each line were almost genetically identical.

In agreement with the history of breeding, the genomes of the three lines are very similar and resemble recombinant inbred strains with C57BL/6J (B6J) and C57BL/10J (B10J) as main backgrounds. After the analysis of the 366 polymorphic SNPs present in the MiniMUGA array it was estimated that the percentage backgrounds were 55.6% C57BL/10J and 44.4% C57BL/6J for line LE-Tg; 59.6% C57BL/10J and 40.4% C57BL/6J for line SE-Tg; and 69.1% C57BL/10J and 31.5% C57BL/6J for line PE-Tg. Using a special function from a proprietary spreadsheet developed by Transnetyx, we could rule out the presence of “diagnostic SNPs” from C57BL/6N (NIH substrain) in the three Tg lines. With the same tool we could also rule out traces of DBA/2 alleles in line 0301. However, we estimated the presence of a very small (~1%) proportion of SWR/J genome in lines SE-Tg and PE-Tg.

WT C57BL/6J (B6J, strain # 000664), C57BL/10J (B10J, strain # 000665) and SWR/J (strain # 000689) mice were purchased from The Jackson Laboratories.

All mice were housed under specific pathogen-free and temperature-controlled (25°C) conditions with a 12 h light-dark cycle. *Ex vivo* experiments were carried out using 10–12-week-old male mice. For *in vivo* experiments, female or male mice were used at 8 weeks of age. Mice were administered intraperitoneal (IP) injections containing 100 μl of either PBS, or recombinant mouse IFN-γ (10 µg per injection) twice a week. All animal experiments were approved by the Institutional Animal Care & Use Committee (IACUC) at the University of Michigan.

### Reagents

All materials used, along with vendor names and catalog numbers are listed in Supplementary Table 5.

### Cell culture

THP-1 cells were cultured in RPMI 1640 culture medium (Invitrogen) supplemented with 10% Fetal Bovine Serum (FBS, Corning), 2 mmol/L of glutamine (Gibco) and the antibiotics 100 U/mL penicillin and 100 µg/mL streptomycin (Pen Strep, Gibco), based on a previously described protocol ^66,67^. Briefly, cells were seeded and differentiated using 85 nM Phorbol 12-Myristate 13-Acetate (PMA, Sigma) for 3 days, media was replaced, and cells were allowed to rest for 5 additional days. Medium was replaced and cells were subjected to treatments with 100 µg/mL of the peptides 65-79*LE, 65-79*SE or 65-79*PE with or without recombinant human IFN-γ (10 ng/mL).

RAW 264.7 cells were cultured in DMEM culture medium (Gibco) supplemented with 10% FBS (Corning) and 100 U/mL penicillin and 100 µg/mL streptomycin (Pen Strep, Gibco). Cells were plated and treated with 65-79*LE, 65-79*SE or 65-79*PE (100 µg/mL) with or without recombinant mouse IFN-γ (5 ng/mL).

### BMDM differentiation

For primary macrophage experiments, bone marrow cells isolated from femurs and tibias were used to generate bone marrow-derived macrophages (BMDMs), as previously ^20^. For ATP experiments, bone marrow cells were cultured in 6-well plates (2 × 10^6^ cells per well) in α-Minimum Essential Medium (α-MEM), supplemented with 10% (v/v) FBS and 100 U/mL penicillin and 100 μg/mL streptomycin (all from Invitrogen), in the presence of recombinant mouse macrophage colony stimulating factor (M-CSF, 10 ng/mL) for 3 days. Cell culture media were replaced daily. After 3 days, cells were treated with recombinant mouse IFN-γ, (5 ng/mL) for 24 h. For all other functional experiments, bone marrow cells were cultured in DMEM medium, supplemented with 20% (v/v) L929 cell conditioned media, 0.5% (v/v) pyruvate, 10% (v/v) FBS and 100 U/mL penicillin and 100 μg/mL streptomycin (all from Invitrogen) for 3 days, as described previously ^68^. Thereafter, cells were plated for 24 h, followed by treatment with IFN-γ (5 ng/mL) for 48-72 h.

### RNA-Sequencing

Samples for RNA-seq were prepared and processed as previously described ^20^. Briefly, RAW 264.7 cells were treated with 15mer ligands 65-79*LE, 65-79*SE or 65-79*PE (100 μg/mL) with or without IFN-γ (5 ng/mL). Media were refreshed after 48 h, and at 72 h, cells were lysed with Trizol (Invitrogen). Total RNA was isolated using the Direct-Zol^TM^ RNA miniprep kit (Zymo Research) according to manufacturer’s instructions. RNA from THP-1 cells was isolated 72 h after treatment with 15mer ligands 65-79*LE, 65-79*SE or 65-79*PE (100 μg/mL) with or without IFN-γ (10 ng/mL) using the RNeasy mini kit (Qiagen) according to the vendor instructions. Genomic DNA was removed using the Turbo DNA-Free kit (Ambion). All samples used for library generation had an RNA integrity number of 7.5 or higher.

The TruSeq Stranded mRNA Sample Preparation Kit (Illumina) was used for library generation, according to manufacturer’s protocol, using 200ng of total RNA. For each sample, 100bp single-end reads were generated using Illumina’s HiSeq2500v4. Raw read counts were obtained using featureCounts from the Rsubread1.5.0p3 package and Gencode-M12 gene annotations (RAW 264.7) or featureCounts from the Rsubread-1.6.1 package and Gencode28-hg38 gene annotations (THP-1) using only uniquely aligned reads. Data normalization and differential expression analysis were performed using DESeq2-1.16.1 within R-3.4.1 using an adjusted p-value threshold of 0.05. Mouse gene annotations were verified using the MGI database (http://www.Informatics.jax.org/index.shtml) ^69^, only protein encoding genes were considered. Differentially expressed genes with a fold change (FC) of > 1.5 and adjusted *p*-value (Padj) of < 0.05 were used for Gene ontology (GO) and Kyoto Encyclopedia of Genes and Genomes (KEGG) pathway analysis using the DAVID bioinformatics database (version 6.8, https://david.ncifcrf.gov/) ^70^. GO and KEGG terms with a false discovery rate (FDR) of < 5% (< 0.05) were considered significant. Unsupervised clustering of DEGs was visualized using a web server heatmapper (http://www.heatmapper.ca) ^71^ using average linkage and Pearson correlation distance.

### Upstream regulator analysis

Upstream regulator analysis was performed using Advaita Bio’s iPathwayGuide (iPG, Advaita Bioinformatics, MI, https://advaitabio.com/ipathwayguide/) ^72^. All differentially expressed genes were used to predict upstream regulators that are likely to be activated. Upstream regulators with an adjusted p-value of <0.05 were included in the analysis.

### qRT-PCR

cDNA was synthesized from total RNA using High-Capacity cDNA Reverse Transcription Kit (Thermo Fisher) according to the vendor’s protocol. qRT-PCR was performed on a StepOnePlus or a CFX384 Touch (Bio-Rad) Real-Time PCR system, with a total reaction volume of 10 or 20 µl using Fast SYBR^TM^ Green Master Mix (all reagents from Applied Biosystems). The primers used are listed in the KEY RESOURCES table. Analysis was done with StepOne or CFX Maestro Software using the ΔΔCT method.

### Cell death and viability assays

Lactate dehydrogenase (LDH) release from cells was measured using the Cytotoxicity Detection Kit^PLUS^ (Roche), according to the manufacturer’s protocol. Absorbance was measured at 490 nm using a micro plate reader (Bio-Tek, Synergy™ H1). Cytotoxicity was calculated as described ^73^.

Cell viability was evaluated using the methylthiazolyldiphenyl-tetrazolium bromide (MTT) method ^74,75^. Briefly, cells were rinsed with PBS and a 0.5% (w/v) MTT solution was added for 3 h. Next, formazan crystals formed were solubilized with isopropanol (Sigma) and optical density was measured at 540 nm using a micro plate reader (Bio-Tek, Synergy™ H1).

Caspase-3 activity was determined using EnzChek™ Caspase-3 Assay Kit #2 (Thermo Fisher).

### Mitochondrial function assays

Intracellular ATP levels were measured with the ATPlite Luminescence Assay System (PerkinElmer) following the vendor protocol. Mitochondrial membrane potential was determined using the tetramethylrhodamine (TMRE) dye (Invitrogen), following the manufacturer’s protocol. Mitochondrial ROS was measured using MitoSOX Red superoxide indicator (Molecular Probes) following the supplier’s protocol.

### Immunoblots

Cells were collected and lysed with RIPA buffer (Sigma) containing phosphatase inhibitor phosSTOP and protease inhibitors cOmplete mini EDTA-free (Roche). Protein concentration was quantified using *RC DC*™ Protein Assay Kit (Bio-Rad). Proteins were separated using SDS-PAGE Gels (Thermo Fisher) and transferred to PVDF Western Blotting membranes (Roche). The membranes were blocked with 5% bovine serum albumin (BSA, Sigma). Blots were incubated overnight with primary antibodies listed in the KEY RESOURCES table, followed by 1h incubation with appropriate secondary antibody. Membranes were developed using SuperSignal^TM^ West Pico Plus ECL substrate (Thermo Scientific) or Clarity^TM^ Western ECL Substrate (Bio-Rad). Images were acquired using the Omega Lum™ C Imaging System (Gel Company) or ChemiDoc™ Imaging System (Bio-Rad).

### Immunocytochemistry for ubiquitinylated proteins

Cells were cultured on cover glass and fixed with a 4% paraformaldehyde solution. Fixed cells were subsequently permeabilized with 0.1% triton-X100, blocked in 2% BSA, followed by overnight incubation with mono and polyubiquitinylated conjugates monoclonal antibody (1:200) and 1 h incubation with secondary antibody, labelled with Alexa Fluor 647. Coverslips were mounted onto slides using Prolong Diamond with DAPI (Invitrogen). Cells were imaged using a Nikon E800 Epifluorescence microscope.

### Quantification of DNA damage

DNA damage was quantified using a Comet Assay (Trevigen), based on previously described protocols ^76,77^. The slides were observed by fluorescence microscopy (Cytation 5, GFP filter). A total of 100 cells in each slide were scored according to the length and intensity of the tail in relation to the head as described ^76,77^. Each cell was scored on a scale of 0 to 4 by two independent observers blinded with regards to the samples and the average from both counts was used for calculation ^77^.

### Quantification of cytokines, nitrites, and Serum anti-dsDNA-IgG levels

Cell supernatants were used for quantification of cytokines (IL1-β, IL-6 and TNF-α) by commercial ELISA kits following the protocols provided by the manufacturer (KEY RESOURCES table). To measure nitrite concentrations, cell culture supernatants were collected and immediately subjected to a Griess Reagent assay (Promega), according to the suppliers’ protocol.

Serum anti-dsDNA-IgG levels were measured at week 5, 10, 16, 20 and 24 using an ELISA Kit (Alpha Diagnostic International) according to the manufacturer’s instructions.

### Renal histopathology and immunofluorescence

Renal histopathology was interpreted as described before ^78^. Briefly, mouse kidneys were harvested after 24 weeks of treatment, fixed in formalin, and 3 µm sections were stained with periodic acid–Schiff (PAS) or hematoxylin and eosin (H&E). The kidneys were scored for glomerulitis, extracapillary proliferation and mesangial expansion by a pathologist (E.F.), who was blinded with regards to the experimental groups. Images were taken at a 400 × magnification, with a BX41 microscope (Olympus) and an Olympus DP73 camera using cellSens imaging software.

Complement deposition was quantified by C3 immunofluorescence. Five µm frozen sections were stained with FITC-conjugated anti-C3 (Immunology Consultants Laboratory) for 1 h at 4°C, followed by Hoechst (Invitrogen) counterstaining. Images were captured by a Nikon E800 Epifluorescence and Brightfield microscope (Nikon, Melville, NY) using a 10x objective. Glomerular C3 staining was scored on a 0-3 basis in a blinded manner in at least eight glomeruli per section. Positive controls were identically processed using frozen kidney sections from 28 weeks old female New Zealand Mixed (NZM) 2328 mice.

### Statistical analysis

Unless otherwise specified, data are presented as mean value ± standard error of the mean (SEM). All statistical analyses were performed using the Graph Pad Prism 8.0 statistical software (Graph Pad Software: https://www.graphpad.com/). One-way, or two-way ANOVA were used for analysis of normally distributed *in vitro* data. The statistical significance of the differences of anti-dsDNA IgG (Figs. 7f and 7h) levels and albumin-to-creatinine ratios (ACR) (Fig. 7i) among the different *in vivo* mouse treatment groups was analyzed by a Wilcoxon sign rank test, using the measured levels in PBS-treated LE-Tg, and the measured levels in IFN-γ-treated SE-Tg mice as the theoretical medians for statistical analysis compared to the measured levels in IFN-γ-treated LE-Tg mice, under the assumption that treatment of LE-Tg with IFN-γ induces higher levels. The significance of the differences in renal histopathology scores among the different mouse treatment groups (Fig. 7j) was analyzed by a Wilcoxon sign rank test as well, using a theoretical median of zero under the assumption that the various treatments do not induce any glomerular pathology. For all experiments, the statistical test used and the number of replicates are indicated in the figure legends. *p*-values < 0.05 were considered statistically significant.

### Data Availability

RNA-seq datasets generated in this manuscript were deposited in the NCBI Gene Expression Omnibus and are accessible through GEO accession number GSE173877. Source data are provided with this paper. Additional data supporting the findings of this study are available from the Lead Contact upon request.

## ACKNOWLEDGEMENTS

We thank Drs. Chela David and Veena Taneja from the Mayo Clinic for providing transgenic mice; Kelsey Rampalski and Claire McCrate-Heath at the University of Michigan for technical assistance; Drs. Terrance O’Hanlon and Ejaz Shamim from the National Institutes of Health, Bethesda, MD, USA, for useful comments on the manuscript. This work was supported by the Extramural Program of the National Institute of Arthritis and Musculoskeletal and Skin Diseases (Grants R01AR059085, R61AR073014, R33AR073014, R01AR074930) to J.H., the National Institute of Environmental Health Sciences Extramural Program (Contract HHSN273201600123P) to J.H., and the Intramural Research Program at the National Institute of Environmental Health Sciences, NIH (Grant ES101074) to F.W.M. B.M.S was supported by Training Grant T32AR07080 from the National Institute of Arthritis and Musculoskeletal and Skin Diseases. The content is solely the responsibility of the authors and does not necessarily represent the official views of the National Institutes of Health.

## SUPPLEMENTAL INFORMATION

Supplementary table 1: DEGs with disease relevance in Model A. Related to Fig. 1

Supplementary table 2: DEGs with disease relevance in Model B. Related to Fig. 2

Supplementary table 3: SLE-relevant DEGs shared between mouse RAW 264.7 and human THP-1 macrophages. Related to Fig. 2

Supplementary table 4: Selected URs for Model A and B. Related to Fig. 1 and 2

Supplementary table 5: Materials and reagents

Supplementary Data File 1: RNA-seq data RAW 264.7 macrophages, Model A

Supplementary Data File 2: RNA-seq data RAW 264.7 macrophages, Model B Supplementary Data File 3: RNA-seq data THP-1 macrophages, Model B

## CONTRIBUTIONS

J.H. conceived the project. J.H., B.M.S., B.K. and V.v.D. designed the cell biology experiments.

J.H., V.v.D., B.M.S. and A.H.S. designed the RNA-seq experiments. J.M.K. and J.H. designed the C3 immunofluorescence experiment. V.v.D. performed RNA-seq bioinformatics analysis. B.M.S., V.v.D., J.C.F., B.K., J.L. and R.A.M.F. performed the experiments and statistical evaluations. F.B. performed mouse genomics analyses. E.A.F. read and scored renal histopathology, J.H. and F.W.M. provided funding. J.H. wrote the paper with input from all co-authors. All authors reviewed the findings and approved the final version of the manuscript.

## SUPPLEMENTARY FIGURE LEGENDS

**Supplementary Fig. 1, Background information and terminologies. Related to Figs. 1, 2, 3 and 7.**

**a,** ‘Top’ (left) and ‘side’ (right) views of a three-dimensional ribbon model of the HLA-DR3 molecule focusing on the cusp region. Green: DRα chain; Yellow: DRβ chain; Red: groove peptide (CLIP). The TAHR polymorphic residues 70-74 are highlighted in cyan blue. Images are based on the known crystal structure of DR3 with the CLIP peptide (1A6A.pdb).

**b,** A list of the *DRB1* alleles studied here along with their respective coded TAHR 65-79 amino acid sequences, designations in this study of the different 15mer synthetic peptides corresponding to these TAHRs, short functional designation of allelic epitopes, and *DRB1* allele-specific transgenic mouse lines used in this study.

**c,** IFN-γ is an obligatory co-factor for epitope-activated transcriptional modulation. qRT-PCR analyses of marker genes in RAW 264.7 macrophages treated with or without 100 μg/mL 65-79*LE (green), 65-79*SE (red) or 65-79*PE (blue) for 72 hours, in the presence (solid-color bars) or absence (light-color bars) of IFN-γ (5 ng/mL). Data represent mean ± SD of 3 independent experiments. One-way ANOVA, * p < 0.05, ** p < 0.01, *** p < 0.001, ****p < 0.0001.

**d,** Allele specificity. Intracellular ATP levels in RAW 264.7 macrophages exposed to epitope-specific 15mer peptides studied here: 65-79*LE (green), 65-79*SE (red), 65-79*PE (blue), along with two additional, control synthetic 15mer allelic peptides: 65-79*0403 (corresponding to allele *DRB1*04:03*) and 65-79*1501 (corresponding to allele *DRB1*15:01*) in the presence of IFN-γ.

Data represents mean ± SD from 3 replicates. One-way ANOVA, ** p < 0.01, *** p < 0.001, ****p < 0.0001.

**e,** Dose-response curves. Intracellular ATP levels in RAW 264.7 macrophages treated with various doses of 65-79*LE, 65-79*SE and 65-79*PE and IFN-γ (5ng/mL). Data represent mean ± SEM of 3 biological replicates.

**Supplementary Fig. 2, LE-activated transcriptional modulation in THP-1 and RAW 264.7 macrophages (Model B). Related to Fig. 2.**

**a,** Venn diagrams showing DEG comparisons in 65-79*LE-, 65-79*SE- and 65-79*PE-stimulated THP-1 macrophages.

**b**,**c**, Notable enriched GO-BP terms for 65-79*LE-upregulated DEGs in RAW 264.7 **b,** and THP-1

**c,** macrophages.

**d**,**e**, Regulatory networks comparing the interactions, mechanisms and regulator modes for STAT1 in THP-1 macrophages activated by 65-79*LE **d,** versus 65-79*SE **e,**.

**f,** Regulatory network legend (iPG).

**Supplementary Fig. 3: Meta-analysis of Model A and Model B. Related to Figs. 1 and 2.**

**a,** Venn diagrams showing meta-comparisons of DEGs in Model A (darker color shades) and Model B (lighter color shades) for 65-79*LE (Green), 65-79*SE (Red) and 65-79*PE (Blue).

**b**, Venn diagrams showing the extent of overlaps of upregulated or downregulated DEGs by 65-79*LE (Green), 65-79*SE (Red) and 65-79*PE (Blue) that found to be shared in **a** between Model A and Model B.

**c, d**, Unique GO-BP terms enrichment among upregulated DEGs that are shared between Model A and Model B. Green, 65-79*LE; Red, 65-79*LE.

**e**, KEGG pathway enrichment among upregulated DEGs that are shared between Model A and Model B Green, 65-79*LE; Red, 65-79*LE.

**Supplementary Fig. 4: LE-triggered ER stress and mitochondrial dysfunction in mouse RAW 264.7 macrophages. Related to Fig. 3.**

**a,** Representative immunoblots of ER stress markers pIRE1-α, GRP78 and CHOP in RAW 264.7 macrophages exposed to different allelic epitopes in the presence of IFN-γ.

**b,** qRT-PCR analysis of *Grp78* expression in RAW 264.7 macrophages exposed to different allelic epitopes in the presence of IFN-γ.

**c**, **d**, Immunoblots of p62 (**c**) and poly-ubiquitinated proteins (**d**) in RAW 264.7 macrophages stimulated by different allelic epitopes in the presence of IFN-γ.

**e,** Representative TMRE immunocytochemistry images of allelic epitope-exposed RAW 264.7 macrophages in the presence of IFN-γ. Scale bar=100 μm.

**f,** Mitochondrial ROS in RAW 264.7 macrophages exposed to allelic epitopes in the presence of IFN-γ and measured by MitoSOX.

**g,** Representative MitoSOX immunocytochemistry images of allelic epitope-exposed THP-1 macrophages in the presence of IFN-γ. Scale bar=100 μm.

Blots (**a**, **c**, **d**) are representative of 3 independent experiments. Bar graphs (**b**, **f**) represent mean ± SEM, one-way ANOVA of repeated measures/Tukey, *p < 0.05; **p < 0.01; ***p < 0.001, ****p < 0.0001.

**Supplementary Fig. 5: The LE triggers cell death and pro-inflammatory cytokine production in mouse RAW 264.7 macrophages and primary BMDMs from WT mouse strains. Related to Fig. 4.**

**a**, Cell death (n=3) and **b,** viability assessed by MTT (n=4) of RAW 264.7 macrophages treated with different allelic epitopes.

**c**-**e**, Levels of pro-inflammatory cytokines TNF-α (n=3) **c,** and IL-6 (n=4) **d,** as well as nitrite levels (n=3) **e,** in supernatants of allelic epitope-treated RAW 264.7 macrophage.

**f,** TNF-α and **g,** nitrite supernatant levels in BMDMs derived from control WT mouse strains B6J, B10J and SWR, cultured *ex vivo* in the presence or absence of IFN-γ (5 ng/mL).

Data represent mean ± SEM. One-way (**b-e**), or two-way (**a, f, g**) ANOVA of repeated measures/Tukey. *p < 0.05; **p < 0.01; ***p < 0.001, ****p < 0.0001.

**Supplementary Fig. 6, Additional characterization of LE-activated cell death aberrations. Related to Fig. 5 and Fig. 6.**

**a,** The pan-caspase inhibitor ZVAD-FMK (10 µM) does not hinder 65-79*LE-activated cell death in RAW 264.7 macrophages (n=3).

**b,** Rapamycin (50 nM) has an allele-nonspecific inhibitory effect on 65-79*LE-activated cell death in RAW 264.7 macrophages (n=3).

**c, d,** LC3 II/ LC3 I ratio in allelic epitope-treated RAW 264.7 **c,** (n=4) and THP-1 **d,** (n=3) macrophages.

**e,** Immunoblot of Beclin1 in different epitope-treated RAW 264.7 macrophages. A representative blot, one of 3 independent experiments.

**f,** Necrostatin-1 (50 µM) shows a modest, allele-nonspecific inhibitory effect on LE-activated levels of TNF-α in RAW 264.7 macrophage supernatants (n=5).

**g,** A TNF-α inhibitor [6,7-Dimethyl-3-((methyl-(2-(methyl-(1-(3-trifluoromethyl-phenyl)-1H-indol-3-ylmethyl)-amino)-ethyl)-amino)-methyl)-chromen-4-one] (10 μM) blocks 65-79*LE-activated cell death in an allele-nonspecific fashion.

**h,** ER stress pathways and their respective inhibitors.

**i**, ATF6α pathway signaling blocker, Ceapin-A7 (10 μM) shows allele-specific effect on intracellular levels of ATP in RAW 264.7 macrophage supernatants (n=3).

**j**, **k,** PERK (EIF2AK3) pathway inhibitor, AMG PERK 44 (25 μM) and ER transmembrane protein IRE1 inhibitor, 4μ8C (10 μM) do not rescue intracellular ATP levels in RAW 264.7 macrophages (n=3).

In (**a**, **b**, **f**-**k**), solid-color and dotted-color bars represent, respectively, absence or presence of inhibitors.

Data represent mean ± SEM. Two-way ANOVA of repeated measures/Tukey. *p < 0.05; **p < 0.01; ***p < 0.001, ****p < 0.0001.

**Supplementary Fig. 7: LE-effects in control mice BMDMs Related to Fig. 7.**

**a**, **b**, Immunoblots of ER stress (pIRE1-α, CHOP, GRP78) and proteasomal degradation (p62) markers (**a**) and poly-ubiquitinated proteins (**b**) in BMDMs derived from WT control mouse strains B6J, B10J and SWR, cultured *ex vivo* in the presence or absence of IFN-γ (5 ng/mL).

**c**, Intracellular ATP in BMDMs derived from control mice expressing B6J, B10J and SWR, cultured *ex vivo* in the presence or absence of IFN-γ (5 ng/mL).

Data represent mean ± SEM. One-way ANOVA (**c**).

**Supplementary Fig. 8: IFN-I gene expression levels in BMDMs from Tg mice. Related to Fig. 7.**

qRT-PCR analysis of the expression levels of salient IFN-I genes in BMDMs derived from transgenic mice (n=6) expressing physiologically folded HLA-DRβ molecules coded by *DRB1*03:01* (LE-Tg) or *DRB1*04:01* (SE-Tg) and cultured *ex vivo* for 24h in the presence or absence of IFN-γ (5 ng/mL). Results represent gene expression relative to LE-Tg without IFN-γ treatment.

Data represent mean ± SEM, one-way ANOVA of repeated measures/Tukey, *p < 0.05; **p < 0.01; ***p < 0.001, ****p < 0.0001.

