## Supplemental Figures for "The lupus susceptibility allele *DRB1*03:01* encodes a disease-driving epitope"

**a**

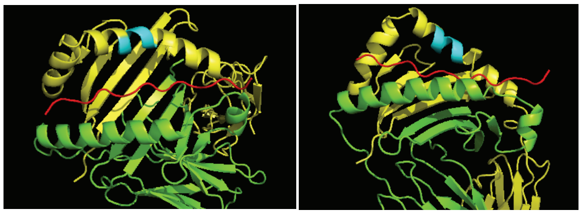

**b**

| Allele | TAHR sequence | Peptide | Epitope | Mouse line |
|---|---|---|---|---|
| *HLA-DRB1*03:01* | 65-KDLLEQKRGRVDNYC-79 | 65-79*LE | LE | LE-Tg |
| *HLA-DRB1*04:01* | 65-KDLLEQKRAAVDTYC-79 | 65-79*SE | SE | SE-Tg |
| *HLA-DRB1*04:02* | 65-KDILEDERAAVDTYC-79 | 65-79*PE | PE | PE-Tg |

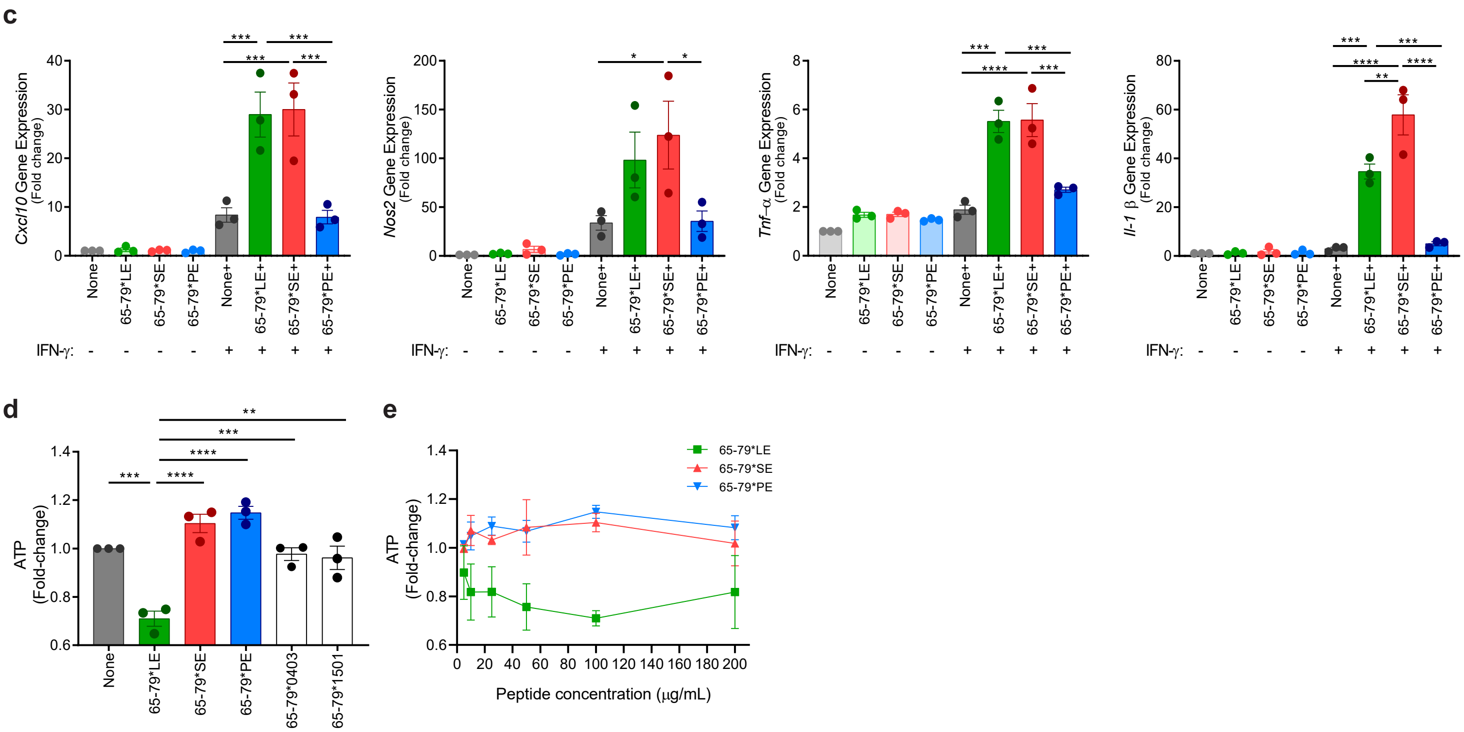

**Supplementary Fig. 1: Background information and terminologies. Related to Figs. 1, 2, 3 and 7.**

**a**, 'Top' (left) and 'side' (right) views of a three-dimensional ribbon model of the HLA-DR3 molecule focusing on the cusp region. Green: DRα chain; Yellow: DRβ chain; Red: groove peptide (CLIP). The TAHR polymorphic residues 70-74 are highlighted in cyan blue. Images are based on the known crystal structure of DR3 with the CLIP peptide (1A6A.pdb).
**b**, A list of the *DRB1* alleles studied here along with their respective coded TAHR 65-79 amino acid sequences, designations in this study of the different 15mer synthetic peptides corresponding to these TAHRs, short functional designation of allelic epitopes, and *DRB1* allele-specific transgenic mouse lines used in this study.
**c**, IFN-γ is an obligatory co-factor for epitope-activated transcriptional modulation. qRT-PCR analyses of marker genes in RAW 264.7 macrophages treated with or without 100 µg/mL 65-79*LE (green), 65-79*SE (red) or 65-79*PE (blue) for 72 hours, in the presence (solid-color bars) or absence (light-color bars) of IFN-γ (5 ng/mL).
**d**, Allele specificity. Intracellular ATP levels in RAW 264.7 macrophages exposed to epitope-specific 15mer peptides studied here: 65-79*LE (green), 65-79*SE (red), 65-79*PE (blue), along with two additional control synthetic 15mer allelic peptides: 65-79*0403 (corresponding to allele *DRB1*04:03*) and 65-79*1501 (corresponding to allele *DRB1*15:01*) in the presence of IFN-γ.
**e**, Dose-response curves. Intracellular ATP levels in RAW 264.7 macrophages treated with various doses of 65-79*LE, 65-79*SE and 65-79*PE and IFN-γ (5 ng/mL).
Data (**c**-**e**) represent mean ± SD of 3 independent experiments. Two-way (**c**) or One-way (**d**,**e**) ANOVA, * $p < 0.05$, ** $p < 0.01$, *** $p < 0.001$, ****$p < 0.0001$.

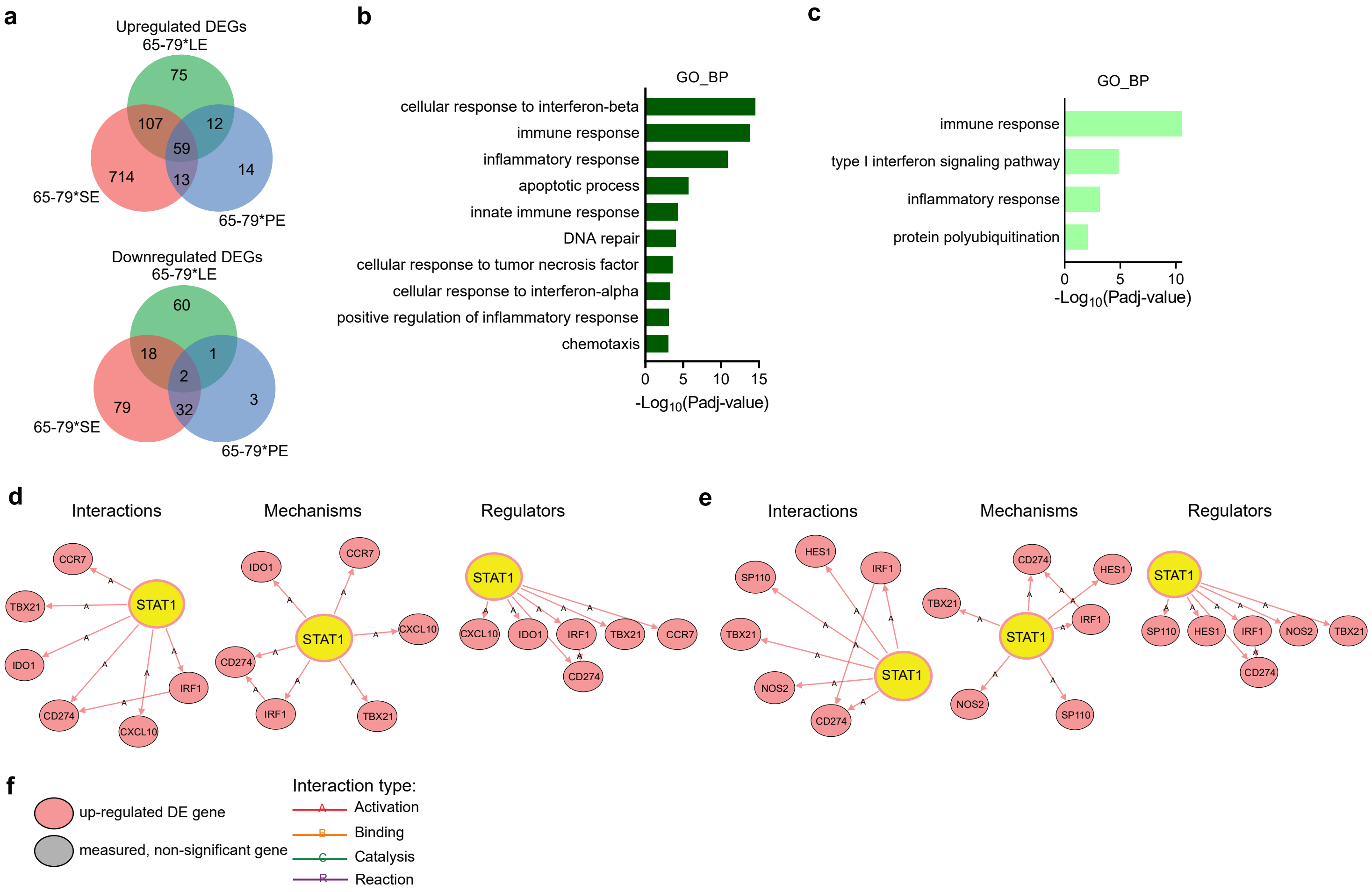

**Supplementary Fig. 2: LE-activated transcriptional modulation in THP-1 and RAW 264.7 macrophages (Model B). Related to Fig. 2.**

**a**, Venn diagrams showing DEG comparisons in 65-79*LE-, 65-79*SE- and 65-79*PE-stimulated THP-1 macrophages.
**b**,**c**, Notable enriched GO-BP terms for 65-79*LE-upregulated DEGs in RAW 264.7 **b**, and THP-1 **c**, macrophages.
**d**,**e**, Regulatory networks comparing the interactions, mechanisms and regulator modes for STAT1 in THP-1 macrophages activated by 65-79*LE **d**, versus 65-79*SE **e**,.
**f**, Regulatory network legend (iPG).

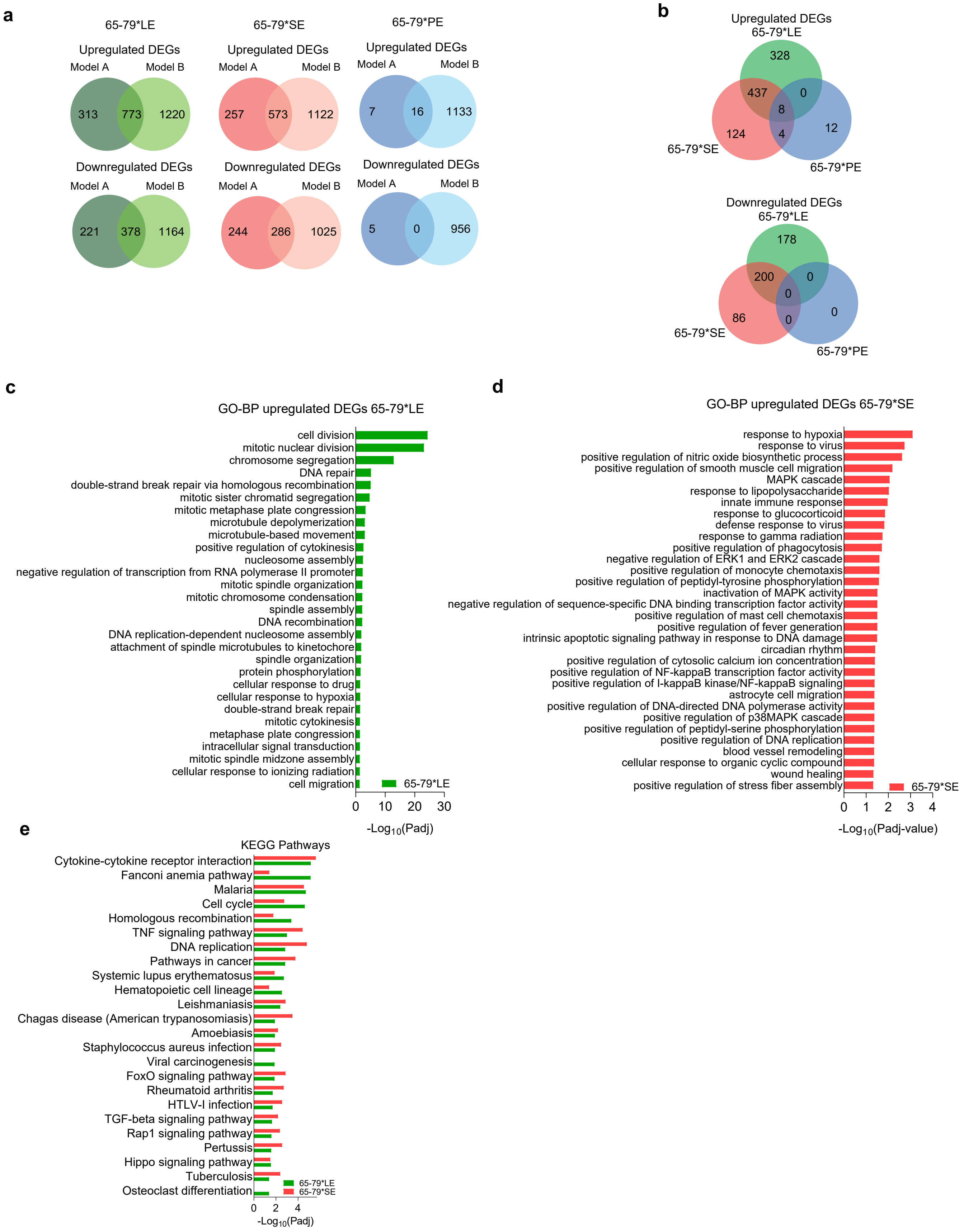

**Supplementary Fig. 3: Background information and terminologies. Related to Figs. 1 and 2.**

**a**, Venn diagrams showing comparison of DEGs in Model A and Model B for 65-79*LE, 65-79*SE and 65-79*PE.
**b**, Venn diagrams showing comparison of DEGs that are shared between Model A and Model B for 65-79*LE, 65-79*SE and 65-79*PE.
**c, d**, Unique GO-BP terms enriched for by upregulated DEGs similar between Model A and Model B 65-79*LE and 65-79*SE.
**e**, KEGG pathway enrichment for upregulated DEGs shared between Model A and Model B for 65-79*LE and 65-79*SE.

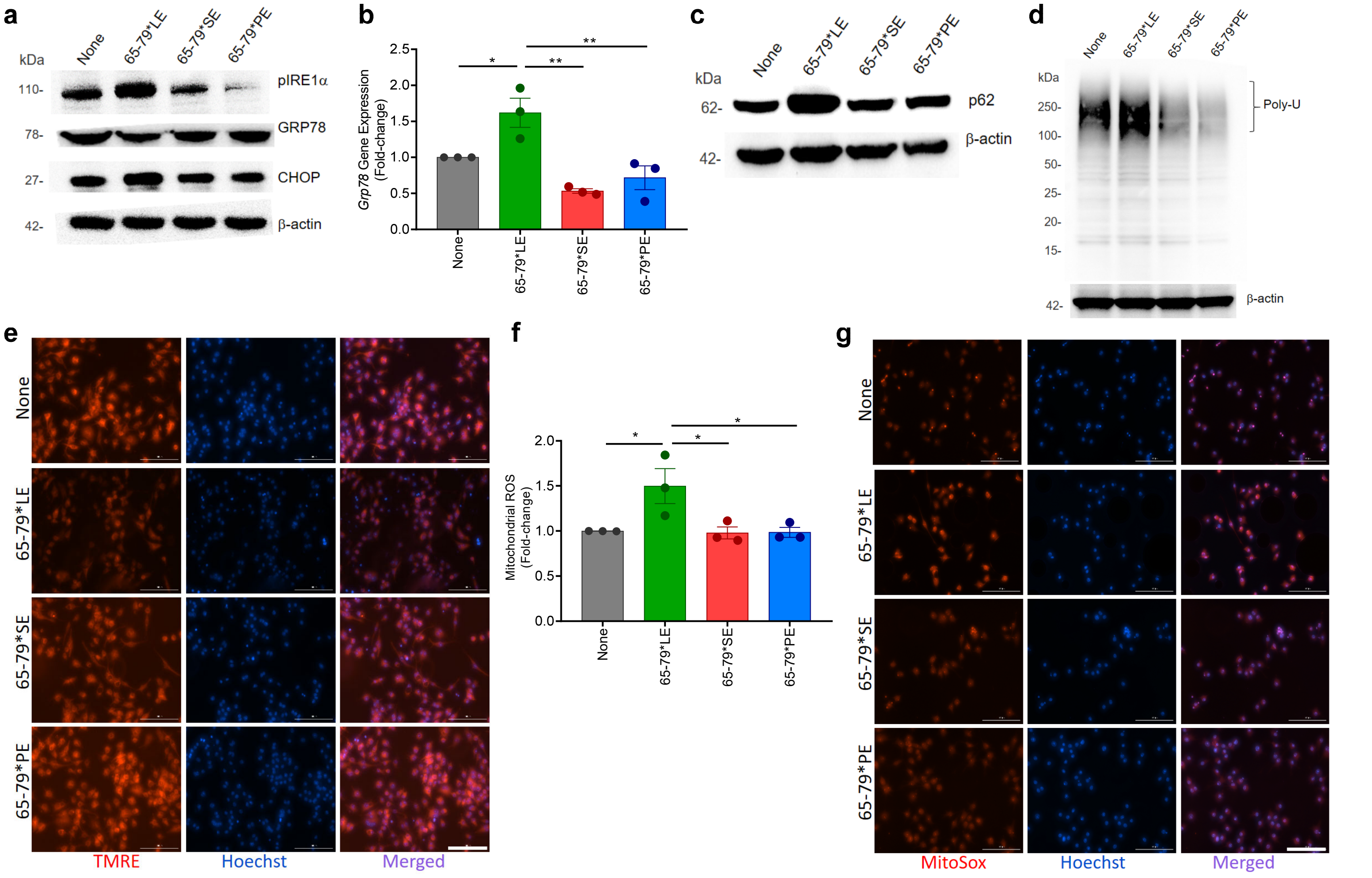

**Supplementary Fig. 4: LE-triggered ER stress and mitochondrial dysfunction in mouse RAW 264.7 macrophages. Related to Fig. 3.**

**a**, Representative immunoblots of ER stress markers pIRE1-α, GRP78 and CHOP in RAW 264.7 macrophages exposed to different allelic epitopes in the presence of IFN-γ.
**b**, qRT-PCR analysis of *Grp78* expression in RAW 264.7 macrophages exposed to different allelic epitopes in the presence of IFN-γ.
**c**, **d**, Immunoblots of p62 **c**, and poly-ubiquitinated proteins **d**, in RAW 264.7 macrophages stimulated by different allelic epitopes in the presence of IFN-γ.
**e**, Representative TMRE immunocytochemistry images of allelic epitope-exposed RAW 264.7 macrophages in the presence of IFN-γ. Scale bar=100 μm.
**f**, Mitochondrial ROS in RAW 264.7 macrophages exposed to allelic epitopes in the presence of IFN-γ and measured by MitoSOX.
**g**, Representative MitoSOX immunocytochemistry images of allelic epitope-exposed THP-1 macrophages in the presence of IFN-γ. Scale bar=100 μm.
Blots (**a**, **c**, **d**) are representative of 3 independent experiments. Bar graphs (**b**, **f**) represent mean ± SEM, one-way ANOVA of repeated measures/Tukey, *p < 0.05; **p < 0.01; ***p < 0.001, ****p < 0.0001.

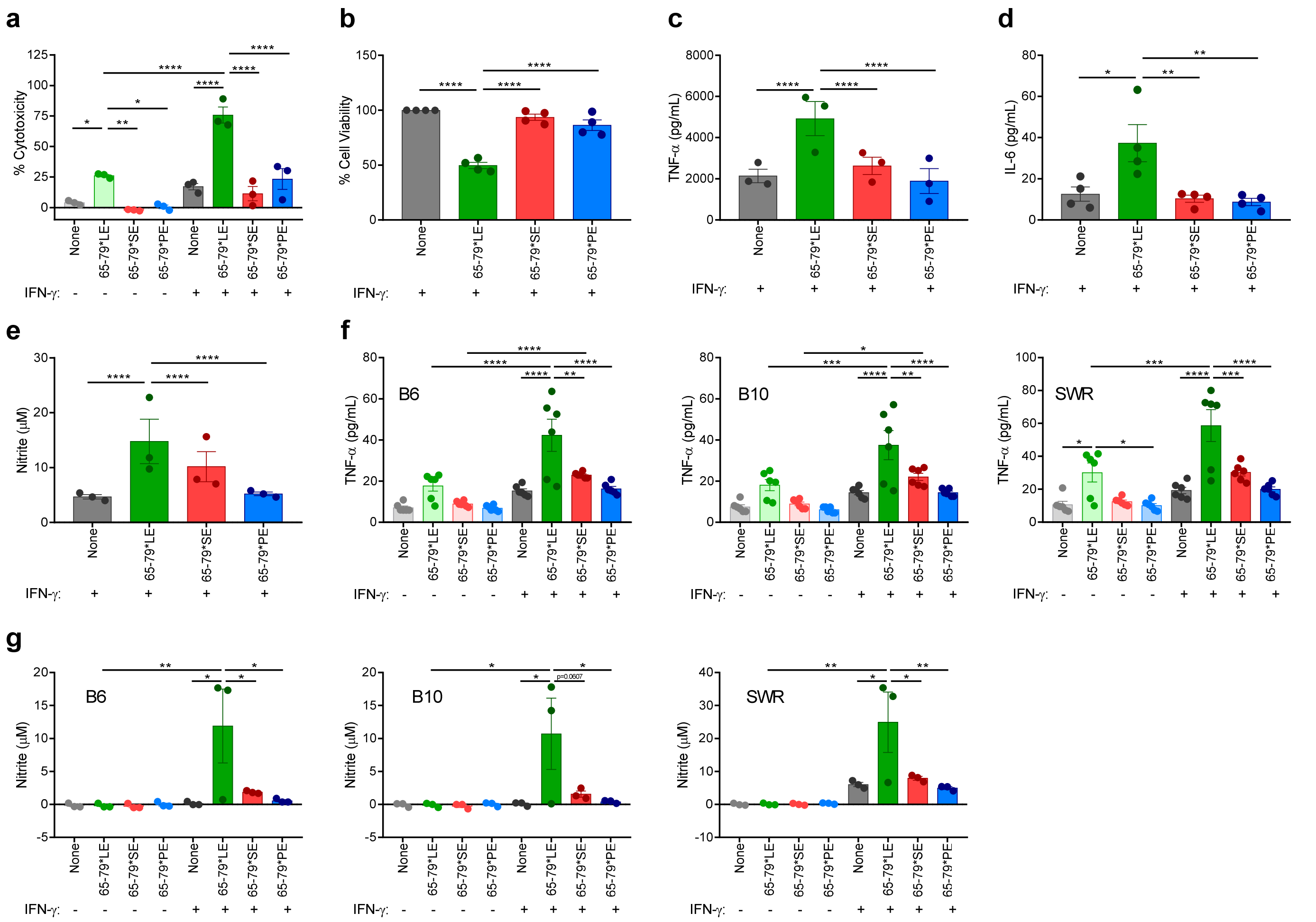

**Supplementary Fig. 5: The LE triggers cell death and pro-inflammatory cytokine production in mouse RAW 264.7 macrophages and primary BMDMs from WT mouse strains. Related to Fig. 4.**

**a**, Cell death (n=3) and **b**, viability assessed by MTT (n=4) of RAW 264.7 macrophages treated with different allelic epitopes. **c**-**e**, Levels of pro-inflammatory cytokines TNF-α (n=3) c, and IL-6 (n=4) **d**, as well as nitrite levels (n=3) **e**, in supernatants of allelic epitope-treated RAW 264.7 macrophage.
Supernatant levels of **f**, TNF-α and **g**, nitrite, in BMDMs derived from control WT mouse strains B6J, B10J and SWR, cultured *ex vivo* in the presence or absence of IFN-γ (5 ng/mL).
Data represent mean ± SEM. One-way (**b**-**e**), or two-way (**a**, **f**, **g**) ANOVA of repeated measures/Tukey. $^*p < 0.05$; $^{**}p < 0.01$; $^{***}p < 0.001$, $^{****}p < 0.0001$.

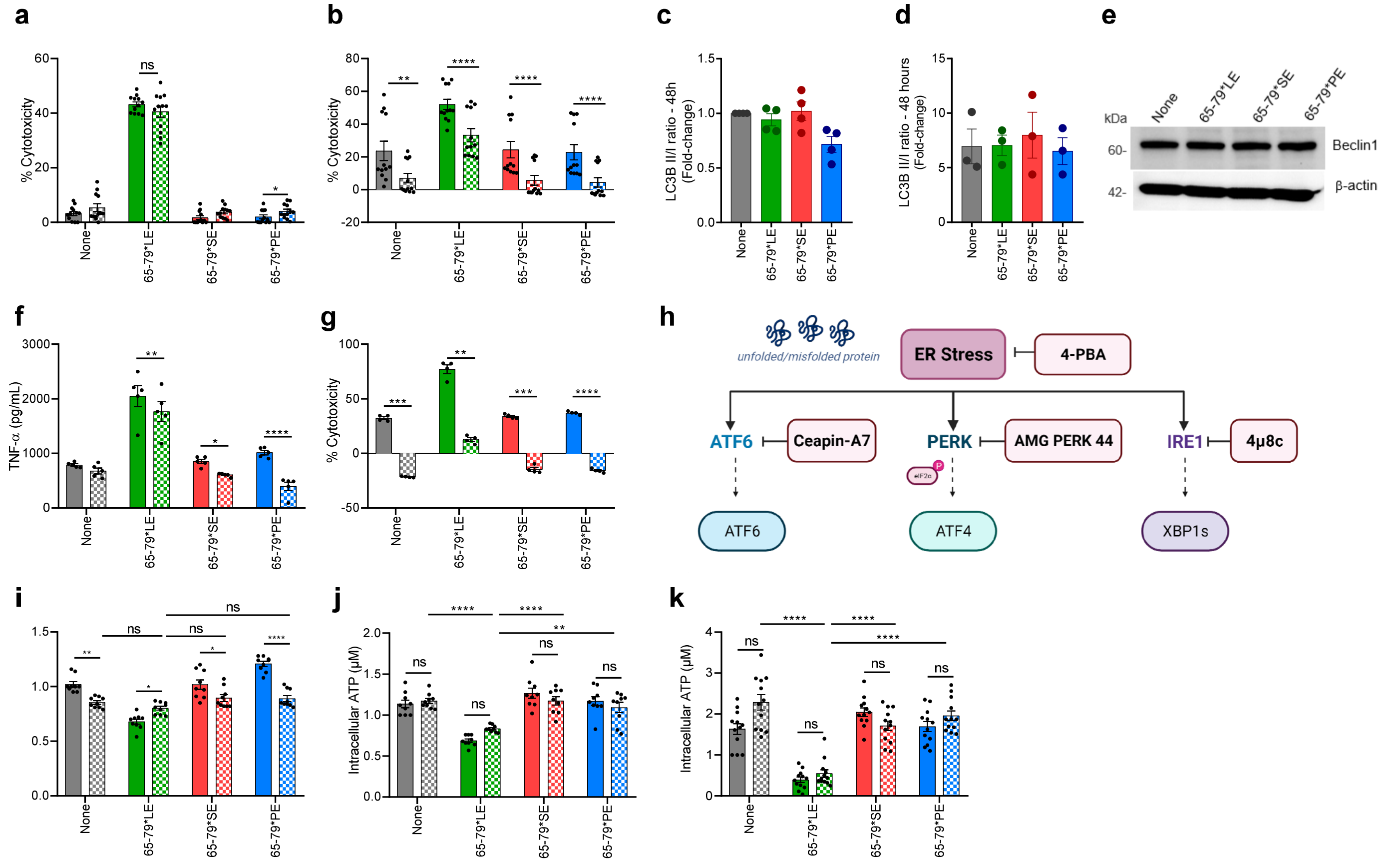

**Supplementary Fig. 6: Additional characterization of LE-activated cell death aberrations. Related to Fig. 5 and Fig. 6.**

**a**, The pan-caspase inhibitor ZVAD-FMK (10 $\mu$M) does not hinder 65-79*LE-activated cell death in RAW 264.7 macrophages (n=3).
**b**, Rapamycin (50 nM) has an allele-nonspecific inhibitory effect on 65-79*LE-activated cell death in RAW 264.7 macrophages (n=3).
**c**, **d**, LC3 II/ LC3 I ratio in allelic epitope-treated RAW 264.7 **c**, (n=4) and THP-1 **d**, (n=3) macrophages.
**e**, Immunoblot of Beclin1 in different epitope-treated RAW 264.7 macrophages. A representative blot, one of 3 independent experiments.
**f**, Necrostatin-1 (50 $\mu$M) shows a modest, allele-nonspecific inhibitory effect on LE-activated levels of TNF-$\alpha$ in RAW 264.7 macrophage supernatants (n=5).
**g**, A TNF-$\alpha$ inhibitor [6,7-Dimethyl-3-((methyl-(2-(methyl-(1-(3-trifluoromethyl-phenyl)-1H-indol-3-ylmethyl)-amino)-ethyl)-amino)-methyl)-chromen-4-one] (10 $\mu$M) blocks 65-79*LE-activated cell death in an allele-nonspecific fashion.
**h**, ER stress pathways and their respective inhibitors.
**i**, ATF6$\alpha$ pathway signaling blocker, Ceapin-A7 (10 $\mu$M) shows allele-specific effect on intracellular levels of ATP in RAW 264.7 macrophage supernatants (n=3).
**j**, **k**, PERK (EIF2AK3) pathway inhibitor, AMG PERK 44 (25 $\mu$M) and ER transmembrane protein IRE1 inhibitor, 4$\mu$8C (10 $\mu$M) do not rescue intracellular ATP levels in RAW 264.7 macrophages (n=3).
In (**a**, **b**, **f-k**), solid-color and dotted-color bars represent, respectively, absence or presence of inhibitors.
Data represent mean ± SEM. Two-way ANOVA of repeated measures/Tukey. *$p < 0.05$; **$p < 0.01$; ***$p < 0.001$, ****$p < 0.0001$.

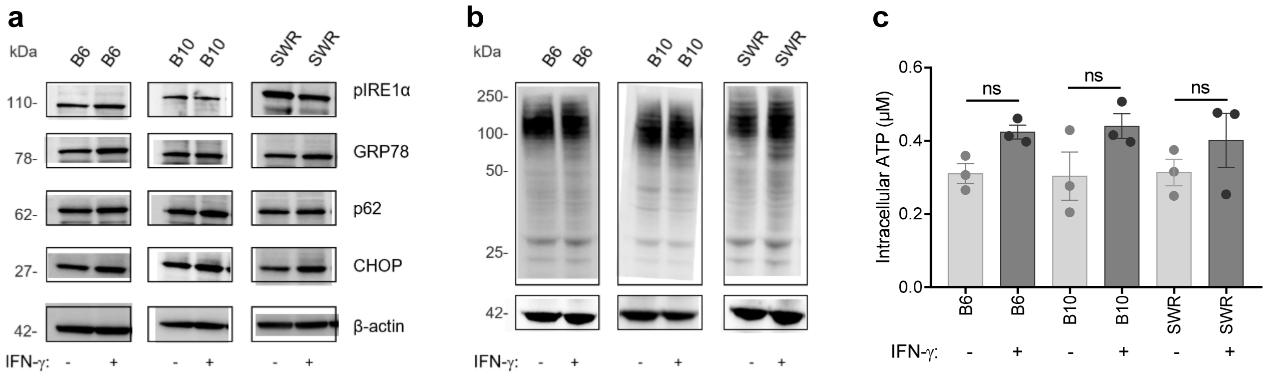

**Supplementary Fig. 7: LE-effects in control mice BMDMs Related to Fig. 7.**

**a**, **b**, Immunoblots of ER stress (pIRE1-α, CHOP, GRP78) and proteasomal degradation (p62) markers **a**, and poly-ubiquitinated proteins **b**, in BMDMs derived from WT control mouse strains B6J, B10J and SWR, cultured ex vivo in the presence or absence of IFN-γ (5 ng/mL).

c, Intracellular ATP in BMDMs derived from control mice B6J, B10J and SWR, cultured ex vivo in the presence or absence of IFN-γ (5 ng/mL).

Data represent mean ± SEM. One-way ANOVA (**c**).

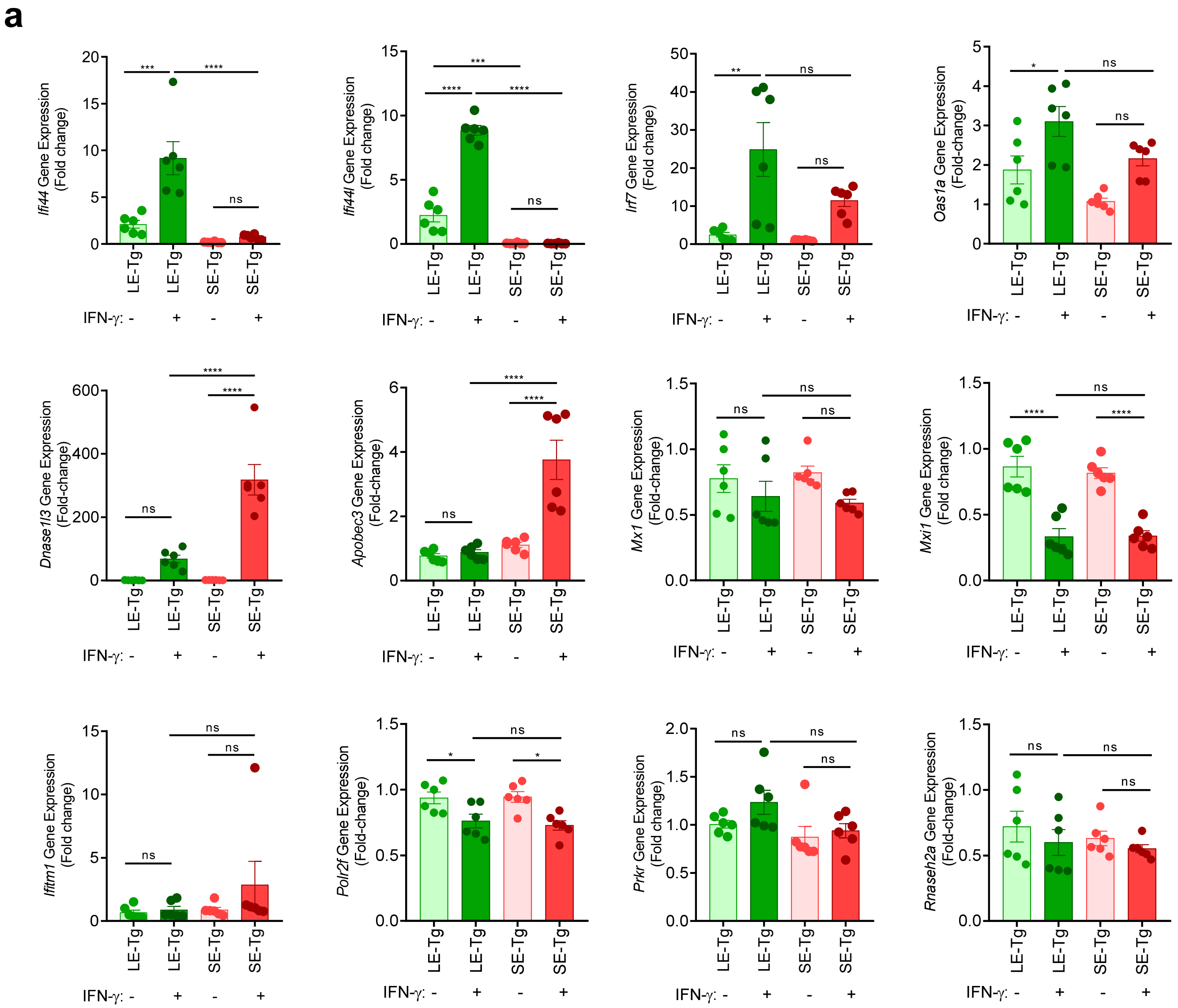

**Supplementary Fig. 8: IFN-I gene expression levels in BMDMs from Tg mice. Related to Fig. 7**

**a**, qRT-PCR analysis of the expression levels of salient IFN-I genes in BMDMs derived from transgenic mice (n=6) expressing physiologically folded HLA-DRβ molecules coded by *DRB1*03:01* (LE-Tg) or *DRB1*04:01* (SE-Tg) and cultured *ex vivo* for 24h in the presence or absence of IFN-γ (5 ng/mL). Results represent gene expression relative to LE-Tg without IFN-γ treatment. Data represent mean ± SEM, one-way ANOVA of repeated measures/Tukey, $^{*}p < 0.05$; $^{**}p < 0.01$; $^{***}p < 0.001$, $^{****}p < 0.0001$.

#### Supplementary Table 1: DEGs with disease relevance in Model A, related to Figure 1

##### Supplementary Table 1A: Notable SLE-relevant genes modulated by 65-79*LE

| Upregulated | FC | Padj | Function |
|---|---|---|---|
| *Ccl5* | 3.97 | $2.07 \times 10^{-66}$ | Upregulated in lupus mice and LN urine |
| *Cxcl2* | 4.62 | $3.19 \times 10^{-63}$ | Upregulated in LN; biomarker for active SLE |
| *Pim1* | 3.35 | $9.42 \times 10^{-55}$ | Therapeutic target in LN; STAT-induced signaling |
| *Lig1* | 2.68 | $4.08 \times 10^{-49}$ | DNA damage repair |
| *Myc* | 4.68 | $3.32 \times 10^{-46}$ | Upregulated in SLE; regulates apoptosis |
| *Ccl3* | 3.32 | $1.98 \times 10^{-42}$ | Autoantibody target is SLE; increased in discoid lupus |
| *Dusp1* | 3.82 | $2.79 \times 10^{-39}$ | Regulate T cell-mediated autoimmune responses |
| *Slamf7* | 2.05 | $3.29 \times 10^{-34}$ | Altered expression in SLE |
| *Tnfrsf9* | 4.07 | $1.84 \times 10^{-33}$ | CD137, expressed on activated T cells; associated with enhanced IFN-$\gamma$ production |
| *Zfp36* | 2.03 | $5.95 \times 10^{-33}$ | Regulates proinflammatory and immune responses; regulates necroptosis |
| *Brca2* | 2.35 | $1.39 \times 10^{-30}$ | DNA damage repair |
| *Chek1* | 2.82 | $2.16 \times 10^{-30}$ | DNA damage response |
| *Timeless* | 2.24 | $8.35 \times 10^{-30}$ | DNA damage repair |
| *Ccl9* | 2.52 | $1.25 \times 10^{-29}$ | Involved in autoimmunity; associated with inflammation |
| *Tnf* | 2.37 | $5.68 \times 10^{-29}$ | Aberrant expression in SLE and autoimmune diseases |
| *Cxcl16* | 2.71 | $9.43 \times 10^{-29}$ | Increased in SLE |
| *Cxcl10* | 3.66 | $4.78 \times 10^{-28}$ | Biomarker of disease severity in LN |
| *Traf1* | 2.25 | $4.10 \times 10^{-27}$ | Polymorphism in SLE |
| *Irf4* | 3.44 | $2.29 \times 10^{-25}$ | Differential expressed in SLE; regulates IFN-driven disease |
| *Dusp4* | 2.57 | $6.58 \times 10^{-25}$ | Overexpressed in T cells of human SLE patients |
| *Smurf1* | 1.62 | $6.35 \times 10^{-25}$ | E3 ubiquitin-protein ligase; plays a role in autoimmunity |
| *Tnfaip3* | 2.61 | $9.86 \times 10^{-25}$ | Deubiquitinating enzyme; susceptibility SNPs in SLE |

| Downregulated | FC | Padj | Function |
|---|---|---|---|
| *Pccb* | 1.76 | $3.91 \times 10^{-56}$ | Plays a role in normal protein processing |
| *Rpn1* | 1.8 | $1.27 \times 10^{-54}$ | Ubiquitin receptor |
| *Scd2* | 2.77 | $2.79 \times 10^{-39}$ | ER enzyme, necessary to prevent ER stress |
| *Erp29* | 1.51 | $2.53 \times 10^{-30}$ | Plays a role in ER stress and the UPR |
| *Hagh* | 1.56 | $3.29 \times 10^{-30}$ | Hydroxyacylglutathione hydrolase, mitochondrial |
| *Pon2* | 1.74 | $1.89 \times 10^{-29}$ | Pro-M2; anti-inflammatory; anti-oxidant; anti-atherogenic |
| *Nucb1* | 1.9 | $5.46 \times 10^{-26}$ | ER protein associated with COX signaling |
| *Marveld1* | 2.15 | $5.90 \times 10^{-25}$ | Reduced Marveld1 increases ROS |
| *Vsir* | 1.69 | $9.08 \times 10^{-25}$ | Reduced expression leads to spontaneous CLE and SLE in mouse |
| *Bcat2* | 1.98 | $3.30 \times 10^{-23}$ | Mitochondrial enzyme |
| *G6pc3* | 1.51 | $6.04 \times 10^{-23}$ | Located in the ER; associated with neutrophil dysfunction/neutropenia |
| *Pdia3* | 1.95 | $1.48 \times 10^{-22}$ | ER protein |
| *Uggt1* | 1.53 | $2.69 \times 10^{-22}$ | Gatekeeper in the ER quality control system |
| *Calr* | 1.89 | $4.93 \times 10^{-22}$ | Defective calreticulin mediated clearance of apoptotic cells in SLE |
| *Arl11* | 1.52 | $1.28 \times 10^{-20}$ | Pro-apoptotic properties |
| *Atg9b* | 2.34 | $2.16 \times 10^{-20}$ | Deficiency potentiates ER stress-associated apoptosis |
| *Aldh18a1* | 1.5 | $2.16 \times 10^{-19}$ | Mitochondrial enzyme |
| *Pnkp* | 1.5 | $3.09 \times 10^{-19}$ | Involved in DNA strand repair |

| | | | |
|---|---|---|---|
| *Cyp27a1* | 1.64 | $3.55 \times 10^{-19}$ | Mitochondrial enzyme |
| *Jmjd8* | 1.74 | $4.38 \times 10^{-19}$ | ER protein |
| *Msmo1* | 1.61 | $4.44 \times 10^{-18}$ | Localized in the ER |
| *Bid* | 1.64 | $6.28 \times 10^{-18}$ | Sensor of cellular stress and DNA damage |
| *Jagn1* | 1.69 | $6.86 \times 10^{-18}$ | Induced in ER stress response |
| *Hsp90b1* | 2.15 | $7.26 \times 10^{-18}$ | GRP94, ER chaperone |
| *Smim14* | 1.89 | $8.65 \times 10^{-18}$ | Possibly plays a role in ER $Ca^{2+}$ homeostasis and ER stress |
| *Prkcsh* | 1.74 | $8.06 \times 10^{-17}$ | Subunit of glucosidase II which is ER located |

**Supplementary Table 1B: Notable RA-relevant genes modulated by 65-79*SE**

| Upregulated | FC | Padj | Function |
|---|---|---|---|
| *Pim1* | 4.00 | $1.64 \times 10^{-72}$ | Regulates pro-inflammatory cytokines in RA |
| *Ccrl2* | 4.46 | $3.95 \times 10^{-59}$ | Upregulated in RA |
| *Pgf* | 13.14 | $9.69 \times 10^{-59}$ | Pro-RA; pro-angiogenesis |
| *Ptpn14* | 7.51 | $8.36 \times 10^{-56}$ | Overexpression in RA FLS |
| *Igf1* | 2.57 | $4.11 \times 10^{-49}$ | Bone modelling |
| *Traf1* | 2.88 | $1.96 \times 10^{-46}$ | RA risk locus |
| *Gpr137b* | 1.83 | $8.20 \times 10^{-45}$ | RA associated locus |
| *Tnfrsf9* | 4.99 | $3.32 \times 10^{-44}$ | CD139, potential RA treatment target |
| *Itgb2* | 1.90 | $1.99 \times 10^{-41}$ | CD18, required for development of inflammatory arthritis |
| *Zfp36* | 2.20 | $3.22 \times 10^{-41}$ | Increased expression in RA; polymorphisms associated with RA; treatment target in RA |
| *Upp1* | 4.35 | $5.71 \times 10^{-38}$ | Increased expression in CIA rat model |
| *Cxcl2* | 3.20 | $6.64 \times 10^{-37}$ | Increased in RA |
| *Cxcl16* | 3.07 | $2.08 \times 10^{-36}$ | Increased expression in RA |
| *Ccl9* | 2.69 | $5.26 \times 10^{-34}$ | Can activate OCs |
| *Tnfaip3* | 2.98 | $7.31 \times 10^{-32}$ | RA risk locus |
| *Ccl22* | 5.93 | $2.67 \times 10^{-31}$ | Potential therapeutic target for RA; expressed in RA synovium |
| *Cd52* | 2.13 | $4.26 \times 10^{-30}$ | Treatment target in RA |
| *Cd83* | 3.10 | $5.37 \times 10^{-29}$ | RA-associated |
| *Dusp4* | 2.71 | $1.03 \times 10^{-27}$ | Pro-angiogenic; pro-Th17 polarization |
| *Pdpn* | 2.20 | $2.26 \times 10^{-26}$ | Implicated in RA; expressed in RA synovium |
| *Adamts1* | 2.15 | $7.38 \times 10^{-26}$ | Involved in angiogenesis |
| *Ninj1* | 1.51 | $8.26 \times 10^{-26}$ | Positively regulates OC development |
| *Rgs1* | 2.91 | $1.37 \times 10^{-25}$ | Involved in inflammation and angiogenesis in RA rats |
| *Rgcc* | 4.44 | $1.59 \times 10^{-25}$ | Dysregulated in RA joint microenvironment |
| *Relb* | 1.81 | $7.56 \times 10^{-25}$ | NF-κB factor associated with RA |
| *Tnip1* | 1.63 | $1.62 \times 10^{-24}$ | Increased expression in RA |

| Downregulated | FC | Padj | Function |
|---|---|---|---|
| *Dock2* | 1.59 | $1.24 \times 10^{-65}$ | Immune regulatory |
| *Scd2* | 3.52 | $7.51 \times 10^{-60}$ | ER enzyme, necessary to prevent ER stress |
| *Sec63* | 1.53 | $1.75 \times 10^{-36}$ | Part of ER protein translocation apparatus |
| *Ero1lb* | 1.68 | $1.02 \times 10^{-32}$ | ER associated markers |
| *Cyb5b* | 1.64 | $9.38 \times 10^{-26}$ | CYB5 mitochondrial isoform |
| *Pon2* | 1.66 | $9.03 \times 10^{-25}$ | Pro-M2; anti-inflammatory; anti-oxidant |

| | | | |
|---|---|---|---|
| *Bet1* | 1.59 | $4.52 \times 10^{-20}$ | Involved in maintenance of mitochondrial functions |
| *Scd1* | 2.51 | $6.50 \times 10^{-20}$ | ER enzyme prevents ER stress |
| *Tmem97* | 1.53 | $5.12 \times 10^{-19}$ | Located in ER membrane; activates mitochondrial superoxide pathway |
| *Irgm2* | 1.56 | $6.12 \times 10^{-18}$ | Inhibits caspase 11 |
| *Vcp* | 1.56 | $9.98 \times 10^{-18}$ | Facilitates polypeptide degradation |
| *Eif2ak3* | 1.62 | $1.20 \times 10^{-17}$ | PERK, part of the UPR |
| *Cd33* | 1.74 | $5.17 \times 10^{-16}$ | Siglec-3, immune-modulatory receptor |
| *Osbp* | 1.5 | $7.20 \times 10^{-16}$ | Regulates ER-Golgi membrane contact formation |
| *Vmac* | 2.24 | $1.14 \times 10^{-15}$ | E3 ubiquitin ligase; involved in STING signaling |
| *Cyp51* | 1.64 | $1.26 \times 10^{-15}$ | Anti-inflammatory |
| *Txndc5* | 1.8 | $1.03 \times 10^{-14}$ | Anti-oxidative stress |
| *Rnf145* | 1.63 | $1.06 \times 10^{-14}$ | Ubiquitin ligase |

### Supplementary Table 2: DEGs with disease relevance in Model B, related to Figure 2

##### Supplementary Table 2A: Notable SLE-relevant genes modulated by 65-79*LE

| Upregulated | FC | Padj | Function |
|---|---|---|---|
| *Cd274* | 9.05 | $1.32 \times 10^{-189}$ | Increased in SLE patients |
| *Cd36* | 10.58 | $9.36 \times 10^{-165}$ | Candidate hub gene in LN; expressed on the majority of monocytes in SLE |
| *Rsad2* | 12.82 | $2.06 \times 10^{-103}$ | Increased expression in SLE |
| *Dio2* | 14.22 | $4.94 \times 10^{-95}$ | Associated with necroptosis |
| *Cxcl10* | 17.98 | $3.86 \times 10^{-66}$ | Biomarker of disease severity in LN |
| *Ifit3* | 7.75 | $4.12 \times 10^{-41}$ | Increased in SLE |
| *Nos2* | 12.87 | $1.05 \times 10^{-39}$ | M1 marker; involved in SLE pathogenesis |
| *Cd69* | 11.51 | $3.00 \times 10^{-37}$ | Can act as a proinflammatory receptor |
| *Myc* | 7.98 | $1.17 \times 10^{-36}$ | Upregulated in SLE |
| *Il27* | 7.72 | $1.15 \times 10^{-34}$ | Augmented in SLE; risk locus for SLE |
| *Rasd2* | 8.9 | $7.76 \times 10^{-20}$ | Main hub gene in LN |
| *Gpr18* | 8.09 | $7.98 \times 10^{-19}$ | Expressed in SLE monocytes |
| *Slc7a2* | 13.77 | $1.41 \times 10^{-17}$ | Required for NO production |
| *Ms4a4c* | 8.25 | $3.18 \times 10^{-17}$ | Upregulated in murine SLE models |
| *Ccl8* | 7.59 | $6.28 \times 10^{-12}$ | Increased in SLE patients with active renal disease |

| Downregulated | FC | Padj | Function |
|---|---|---|---|
| *Dhcr24* | 9.86 | $1.80 \times 10^{-124}$ | Anti-ER stress induced apoptosis |
| *Acaa2* | 3.67 | $2.80 \times 10^{-105}$ | A mitochondrial enzyme; decreased in SLE |
| *Atg9b* | 4.35 | $7.42 \times 10^{-67}$ | Anti-ER stress-associated apoptosis |
| *Alox5* | 4.95 | $3.76 \times 10^{-57}$ | Anti-mitochondrial apoptotic pathway |
| *Nlrp10* | 3.94 | $1.00 \times 10^{-55}$ | Protects against kidney damage; inhibits IL-1β secretion; inhibits NF-κB activation and inhibits apoptosis |
| *Fkbp11* | 4.22 | $3.78 \times 10^{-31}$ | ER stress/UPR gene; anti-inflammation induced apoptosis |
| *Lrrc1* | 4.3 | $7.19 \times 10^{-21}$ | LRRC1-/- cells deregulate Wnt/β-catenin signaling |
| *Cd24a* | 3.24 | $2.40 \times 10^{-14}$ | Triggers caspase-dependent apoptosis; high levels inversely correlated with SLEDAI score |
| *Fhit* | 3.77 | $1.32 \times 10^{-11}$ | Pro-apoptosis |
| *Rab15* | 4.22 | $1.61 \times 10^{-11}$ | Involved in endocytic receptor recycling |
| *Asb10* | 4.22 | $3.68 \times 10^{-10}$ | Functions in ubiquitin-mediated degradation pathways |
| *Map2k6* | 3.41 | $1.07 \times 10^{-7}$ | Pro-survival kinase |

##### Supplementary Table 2B: Notable autoimmune disease-relevant genes modulated by 65-79*PE

| Upregulated | FC | Padj | Function |
|---|---|---|---|
| *Csf1* | 6.54 | $1.91 \times 10^{-32}$ | MCSF; M2 macrophage associated |
| *Bglap2* | 7.19 | $4.93 \times 10^{-30}$ | Anti-bone resorption |
| *Trim30d* | 6.87 | $6.63 \times 10^{-12}$ | Inhibits NF-κB |

| Downregulated | FC | Padj | Function |
|---|---|---|---|
| *Ndrg4* | 5.14 | $1.25 \times 10^{-100}$ | Induced by TNFα through NF-κB |

| | | | |
|---|---|---|---|
| *Ctsk* | 6.57 | $2.22 \times 10^{-63}$ | Osteoclast marker |
| *Mustn1* | 6.1 | $6.77 \times 10^{-47}$ | Over-expressed in arthritis and muscle diseases |
| *Deptor* | 5.65 | $2.63 \times 10^{-40}$ | Involved in mTOR signaling |
| *S100a4* | 4.27 | $1.12 \times 10^{-39}$ | Pro-inflammatory; over-expressed in SLE; biomarker for LN |
| *St6gal1* | 4.03 | $1.42 \times 10^{-38}$ | An SLE disease severity marker |
| *Ndrg2* | 2.93 | $5.10 \times 10^{-29}$ | Pro-inflammatory |
| *Gpr183* | 4.83 | $1.14 \times 10^{-27}$ | Regulates innate and adaptive immunity |
| *Jun* | 3.03 | $4.75 \times 10^{-26}$ | Over-expressed in synovial cells; pro-arthritic in mice |
| *Timp2* | 4.39 | $4.74 \times 10^{-25}$ | Increased in SLE |
| *Cx3cr1* | 8.83 | $1.02 \times 10^{-23}$ | Over-expressed in LN |
| *Tnfrsf9* | 4.17 | $8.61 \times 10^{-20}$ | Autoimmunity-associated locus; over-abundant in RA; therapeutic target in arthritic mice; RA severity locus in African Americans; pro-osteoclastic |
| *Trem2* | 2.93 | $1.62 \times 10^{-19}$ | Regulates OC formation; increased in RA |
| *Rasgrp3* | 3.56 | $8.12 \times 10^{-18}$ | Correlated with SLE disease activity |
| *F10* | 4.54 | $8.40 \times 10^{-18}$ | Factor X, potential key role in RA progression |
| *Slc9b2* | 7.61 | $9.46 \times 10^{-16}$ | Induced by NF-κB |
| *Card14* | 3.62 | $6.10 \times 10^{-15}$ | Induces skin inflammation |
| *Mmp9* | 2.95 | $1.01 \times 10^{-14}$ | Inflammation marker Increased in SLE; SLE-risk locus |
| *Col4a1* | 4.14 | $1.39 \times 10^{-14}$ | Autoantigen in SLE |
| *Clu* | 4.4 | $2.96 \times 10^{-13}$ | Upregulated in inflammatory myopathies |
| *Frat2* | 4.45 | $7.94 \times 10^{-12}$ | Wnt signaling activator |
| *Serpinf1* | 3.94 | $6.83 \times 10^{-10}$ | Mediator of NLRP3 inflammasome |
| *Frat1* | 3.18 | $9.02 \times 10^{-10}$ | Positive regulator of Wnt signaling |
| *F7* | 3.96 | $1.05 \times 10^{-8}$ | Factor VII, plays a role in the pathogenesis of RA; NF-κB and IL-8 activator; pro-angiogenic |
| *Fabp4* | 3.19 | $2.97 \times 10^{-8}$ | Increased levels in RA; pro-angiogenic |
| *Scg2* | 3.52 | $1.12 \times 10^{-7}$ | Pro-angiogenic |

**Supplementary Table 3: SLE relevant DEGs shared between mouse RAW 264.7 and human THP-1 macrophages, related to Figure 2**

| | RAW 264.7 | | THP-1 | | |
|---|---|---|---|---|---|
| | **FC** | **Padj** | **FC** | **Padj** | **Relevant roles/functions** |
| *Psmb9* | 7.89 | $1.22 \times 10^{-174}$ | 1.98 | $4.14 \times 10^{-3}$ | Upregulated in lupus skin |
| *Pim1* | 5.00 | $3.40 \times 10^{-117}$ | 3.64 | $1.50 \times 10^{-2}$ | Proposed therapeutic target for LN |
| *Irf1* | 5.00 | $1.81 \times 10^{-105}$ | 4.16 | $1.46 \times 10^{-3}$ | Induces target gene expression in SLE |
| *Rsad2* | 1.28 | $2.06 \times 10^{-103}$ | 2.61 | $3.98 \times 10^{-2}$ | Upregulated in SLE; IFN signature gene |
| *Stat1* | 6.42 | $1.76 \times 10^{-97}$ | 4.31 | $7.70 \times 10^{-4}$ | Transduces type I and II IFN signaling; overexpressed in SLE |
| *Prdx5* | 3.13 | $7.63 \times 10^{-81}$ | 1.62 | $2.53 \times 10^{-2}$ | Differentially methylated in SLE |
| *Dusp1* | 12.80 | $2.01 \times 10^{-73}$ | 2.60 | $4.96 \times 10^{-2}$ | Regulates T cell-mediated autoimmune responses |
| *Cxcl10* | 1.80 | $3.86 \times 10^{-66}$ | 4.00 | $1.05 \times 10^{-2}$ | Upregulated and proposed as biomarker for active human SLE |
| *Psme1* | 4.61 | $4.35 \times 10^{-60}$ | 1.87 | $3.00 \times 10^{-3}$ | Proteasome activator complex subunit; IFN$\alpha$-inducible gene |
| *Psmb10* | 2.78 | $3.78 \times 10^{-55}$ | 1.82 | $1.29 \times 10^{-2}$ | Immunoproteasome |
| *Cxcl16* | 5.59 | $7.92 \times 10^{-54}$ | 2.50 | $4.66 \times 10^{-2}$ | Increased in SLE; biomarker for disease severity in LN |
| *Stat2* | 3.13 | $1.80 \times 10^{-41}$ | 2.39 | $2.49 \times 10^{-2}$ | Type I IFN-inducible gene; constitutively activated in SLE patients |
| *Cxcl9* | 1.68 | $2.69 \times 10^{-41}$ | 2.43 | $4.67 \times 10^{-11}$ | Overexpressed in cutaneous lupus in correlation with disease activity |
| *Clec2d* | 8.26 | $3.20 \times 10^{-40}$ | 2.03 | $2.54 \times 10^{-2}$ | Induces IFN-$\gamma$ production |
| *Nbn* | 2.21 | $8.73 \times 10^{-35}$ | 1.55 | $5.20 \times 10^{-5}$ | Potential risk locus for SLE |
| *Dusp5* | 18.00 | $6.41 \times 10^{-33}$ | 3.74 | $1.10 \times 10^{-2}$ | Regulates T cell-mediated autoimmune responses |
| *Psme2* | 1.98 | $3.31 \times 10^{-31}$ | 1.81 | $4.80 \times 10^{-2}$ | IFN$\alpha$-inducible gene |
| *Otud1* | 3.73 | $1.95 \times 10^{-25}$ | 1.68 | $4.76 \times 10^{-2}$ | Deubiquinase; mutated in SLE |
| *Nampt* | 1.64 | $3.16 \times 10^{-24}$ | 2.79 | $2.88 \times 10^{-2}$ | Increased in SLE patients |
| *Il15* | 3.09 | $3.91 \times 10^{-21}$ | 3.05 | $2.55 \times 10^{-2}$ | Upregulated in SLE and LN patients |
| *Parp9* | 1.52 | $2.96 \times 10^{-20}$ | 2.23 | $1.07 \times 10^{-2}$ | DNA damage repair; upregulated in SLE; differentially methylated in SLE |
| *Ccl22* | 4.80 | $2.01 \times 10^{-19}$ | 2.36 | $4.76 \times 10^{-2}$ | Risk locus for SLE |
| *Lhfpl2* | 2.78 | $1.59 \times 10^{-13}$ | 1.96 | $1.49 \times 10^{-2}$ | SLE Meta Signature gene |
| *Parp14* | 5.59 | $1.44 \times 10^{-12}$ | 1.93 | $4.44 \times 10^{-2}$ | Acts on M1 polarization downstream of STAT1 |
| *Dtx3l* | 1.69 | $3.99 \times 10^{-8}$ | 1.80 | $1.89 \times 10^{-2}$ | Cooperates with Parp9 in DNA repair; E3 ubiquitin ligase |
| *Scarf1* | 1.70 | $4.63 \times 10^{-5}$ | 2.91 | $3.64 \times 10^{-2}$ | Induces necroptosis (RIPK1/RIPK3-dependent PARP-1 activation) |
| *Tnfsf10* | 2.19 | $5.71 \times 10^{-4}$ | 2.82 | $1.49 \times 10^{-2}$ | Upregulated in LN |
| *Cxcr3* | - 4.19 | $6.17 \times 10^{-21}$ | 2.91 | $1.87 \times 10^{-2}$ | Cxcl9-Cxcl10 receptor; plays a role in murine LN |
| *Svip* | - 2.08 | $9.27 \times 10^{-6}$ | 2.13 | $3.13 \times 10^{-2}$ | Inhibitor of endoplasmic reticulum-associated degradation (ERAD) |

#### Supplementary Table 4: Selected URs for Model A and B, related to Figures 1 and 2

##### Supplementary Table 4A: Top URs activated by 65-79*LE and 65-79*SE in RAW 264.7 macrophages in Model A

| UR | Padj (LE) | Padj (SE) | Relevance |
|---|---|---|---|
| Pclaf | $3.56 \times 10^{-6}$ | $5.86 \times 10^{-4}$ | PCNA; elicits autoimmune responses in SLE; autoantibody target specific for SLE |
| Il1b | $4.02 \times 10^{-6}$ | $8.99 \times 10^{-6}$ | Increased in SLE |
| Egf | $8.94 \times 10^{-6}$ | $1.09 \times 10^{-3}$ | Egfr signaling is involved in LN |
| Egfr | $1.72 \times 10^{-5}$ | $1.43 \times 10^{-3}$ | Egfr signaling is involved in LN |
| Tslp | $1.84 \times 10^{-5}$ | $2.99 \times 10^{-5}$ | Suggested in SLE nephritis |
| Tnf | $2.82 \times 10^{-5}$ | $1.70 \times 10^{-3}$ | Increased in SLE and LN |
| Trp53 | $2.82 \times 10^{-5}$ | $1.70 \times 10^{-3}$ | Increased and autoantibody target in SLE |
| Ccl2 | $4.02 \times 10^{-5}$ | $1.09 \times 10^{-3}$ | Therapeutic target in LN |
| Tlr4 | $4.67 \times 10^{-5}$ | $2.51 \times 10^{-3}$ | Increased in SLE |
| Il33 | $1.15 \times 10^{-4}$ | $4.77 \times 10^{-3}$ | Risk locus in SLE; increased in SLE |
| Tnfsf11 | $2.41 \times 10^{-3}$ | $1.81 \times 10^{-3}$ | RANKL; increased in SLE |
| Csf3 | $2.69 \times 10^{-3}$ | $8.27 \times 10^{-3}$ | Enhances OC activity; increases bone resorption in vivo; increases SLE disease activity and renal involvement |
| Uba52 | $1.71 \times 10^{-4}$ | $1.43 \times 10^{-3}$ | Ubiquitin-60S ribosomal protein L40; source of ubiquitin |
| Rps27a | $5.40 \times 10^{-4}$ | $1.81 \times 10^{-3}$ | Ubiquitin-40S ribosomal protein S27a; source of ubiquitin |
| Ubc | $7.36 \times 10^{-4}$ | $1.39 \times 10^{-2}$ | Poly-ubiquitin C; encodes poly-ubiquitin |
| Mcm9 | $7.80 \times 10^{-4}$ | $2.51 \times 10^{-3}$ | Involved in dsDNA repair |
| Stn1 | $1.98 \times 10^{-3}$ | $1.49 \times 10^{-2}$ | Part of CST complex; contributes to telomere maintenance |
| F2 | $2.69 \times 10^{-3}$ | $8.27 \times 10^{-3}$ | Prothrombin precursor |
| Meiob | $2.69 \times 10^{-3}$ | $8.27 \times 10^{-3}$ | Binds ssDNA during meiosis |
| Clock | $2.73 \times 10^{-3}$ | $8.27 \times 10^{-3}$ | Regulates circadian rhythm |

##### Supplementary Table 4B: Top URs activated by 65-79*LE in THP-1 macrophages in Model B

| Upstream regulator | Padj (LE) | Padj (SE) | Padj (PE) | Relevance to SLE |
|---|---|---|---|---|
| IRF9 | $9.34 \times 10^{-6}$ | | | Type 1 IFN regulated transcription factor |
| IFNG | $1.23 \times 10^{-5}$ | $1.42 \times 10^{-2}$ | $1.01 \times 10^{-5}$ | Mediates SLE pathogenesis; Increased in RA |
| STAT2 | $3.43 \times 10^{-5}$ | | | Increased in SLE |
| STAT1 | $2.99 \times 10^{-3}$ | $2.83 \times 10^{-2}$ | | Increased in SLE; Increased in RA |
| IFNA1 | $3.36 \times 10^{-2}$ | | $3.93 \times 10^{-2}$ | Type 1 IFN, increased in SLE |

#### Supplementary Table 5: Materials and reagents

| Reagent or Resource | Source | Catalog numbers |
|---|---|---|
| **Antibodies (RRID#)** | | |
| Anti-C3, FITC conjugated; (RRID: AB_2891133) | Immunology Consultants Laboratory (Tigard, OR) | Cat# GC3-90F-Z |
| Anti-Mouse CHOP; (RRID:AB_2089254) | Cell Signaling Technologies (Danvers, MA) | Cat# 2895 |
| Anti-Rabbit IRE1 alpha; (RRID:AB_10145203) | Novus Biologicals (Littleton, CO) | Cat# NB100-2323SS |
| Anti-Mouse BiP/Grp78; (RRID:AB_398292) | BD Biosciences (Franklin Lakes, NJ) | Cat# 610979 |
| Anti-Mouse β-actin; (RRID:AB_399901) | BD Biosciences (Franklin Lakes, NJ) | Cat# 612657 |
| Anti-mouse Mono- and polyubiquitinylated conjugates monoclonal (FK2); (RRID:AB_10541840) | Enzo Life science (Farmingdale, NY) | Cat# BML-PW8810 |
| Purified Mouse Anti-RIP; (RRID:AB_397831) | BD Biosciences (Franklin Lakes, NJ) | Cat# 610458 |
| Anti-Rabbit Recombinant Anti-MLKL (phospho S345); (RRID:AB_2687465) | Abcam (Cambridge, United Kingdom) | Cat# ab196436 |
| Anti-Rabbit SQSTM1/p62; (RRID:AB_10624872) | Cell Signaling Technologies (Danvers, MA) | Cat# 5114S |
| Anti-Rabbit LC3B; (RRID:AB_915950) | Cell Signaling Technologies (Danvers, MA) | Cat# 2775S |
| Anti-rabbit IgG, HRP linked; (RRID:AB_2099233) | Cell Signaling Technologies (Danvers, MA) | Cat# 7074S |
| Anti-mouse IgG HRP-linked; (RRID:AB_772210) | GE healthcare Lifesciences (Chicago, IL) | Cat# NA931 |
| Beclin-1 Antibody; (RRID:AB_490837) | Cell Signaling Technologies (Danvers, MA) | Cat# 3738 |
| Goat anti-Mouse IgG (H+L) Cross-Adsorbed Secondary Antibody, Alexa Fluor 647; (RRID:AB_2535804) | Thermo Fisher (Waltham, MA) | Cat# A-21235 |
| IFNAR2 Monoclonal Antibody (MMHAR-2) (RRID:AB_387828) | PBL Assay Science (Piscataway, NJ) | Cat# 21385-1 |
| **Cell Culture** | | |
| Alpha MEM | Gibco (Waltham, MA | Cat# 12561-056 |
| Antibiotics (Pen Strep) | Gibco (Waltham, MA) | Cat# 15140-122 |
| DMEM | Gibco (Waltham, MA) | Cat# 11885-084 |
| DMEM (high glucose) | Gibco (Waltham, MA) | Cat# 11965-092 |
| Fetal Bovine Serum | Corning (Tewksbury, MA) | Cat# 35-015-CV |
| L-Glutamine (200 mM) | Gibco (Waltham, MA) | Cat# 25030-081 |
| RPMI 1640 | Gibco (Waltham, MA) | Cat# 11875-093 |
| Sodium Pyruvate | Gibco (Waltham, MA) | Cat# 11360-070 |
| StemPro™ Accutase™ Cell Dissociation Reagent | Gibco (Waltham, MA) | Cat# A1110501 |

| | | |
|---|---|---|
| **Cells** | | |
| L929 | ATCC (Manassas, VA) | Cat# CCL-1 |
| RAW 264.7 | ATCC (Manassas, VA) | Cat# TIB-71 |
| THP-1 cells | ATCC (Manassas, VA) | Cat# TIB-202 |
| **Peptides** | | |
| 65-79*LE | Bioworld (Dublin, OH) | |
| 65-79*SE | Bioworld (Dublin, OH) | |
| 65-79*PE | Genscript (Piscataway, NJ) | |
| 65-79*1501 | Bioworld (Dublin, OH) | |
| 65-79*0403 | Bioworld (Dublin, OH) | |
| **Chemicals, Reagents, peptides, and Recombinant Proteins** | | |
| 2′,7′-Dichlorofluorescin diacetate (DCFDA) | Sigma (St. Louis, MO) | Cat# D6883 |
| 2x Laemmli Sample Buffer | Bio-Rad (Hercules, CA) | Cat# 1610737 |
| 4μ8C | Sigma (St. Louis, MO) | Cat# SML0949 |
| AMG PERK 44 | Sigma (St. Louis, MO) | Cat# SML3049 |
| Baricitinib | AChemBlock (Hayward, CA) | Cat# G-5743 |
| Bovine Serum Albumin | Sigma (St. Louis, MO) | Cat# A7906 |
| Ceapin-A7 | Sigma (St. Louis, MO) | Cat# SML2330 |
| Clarity™ Western ECL Substrate | Bio-Rad (Hercules, CA) | Cat# 170-5060 |
| cOmplete mini EDTA-free | Roche (Indianapolis, IN) | Cat# 11836170001 |
| Hoechst 33342 | Invitrogen (Waltham, MA) | Cat# H3570 |
| Methylthiazolyldiphenyl-tetrazolium bromide (MTT) | Sigma (St. Louis, MO) | Cat# M2128 |
| MitoSOX Red | Invitrogen (Waltham, MA) | Cat# M36008 |
| Necrostatin-1 (Nec-1) | Enzo Life science (Farmingdale, NY) | Cat# BML-AP309-0020 |
| Necrosulfonamide (NSA) | TOCRIS (Bristol, UK) | Cat#5025 |
| Novex™ 4-20% Tris-Glycine gels | Invitrogen (Waltham, MA) | Cat# XP04200BOX |
| Novex™ WedgeWell 16% Tris-Glycine gels | Invitrogen (Waltham, MA) | Cat# XP0016BOX |
| NuPAGE™ LDS Sample Buffer (4x) | Invitrogen (Waltham, MA) | Cat# NP0007 |
| NuPAGE™ Sample Reducing agent (10x) | Invitrogen (Waltham, MA) | Cat# NP0009 |
| NuPAGE™ 10% Bis-Tris gel | Invitrogen (Waltham, MA) | Cat# NP0301BOX |
| PBS (Phosphate buffered saline) | Gibco (Waltham, MA) | Cat# 10010-023 |
| Phorbol 12-myristate 13-acetate (PMA) | Sigma (St. Louis, MO) | Cat# P8139 |
| phosSTOP | Roche (Indianapolis, IN) | Cat# 04906845001 |
| Pluronic™ F-127 | Invitrogen (Waltham, MA) | Cat# P3000MP |
| ProLong™ Diamond Antifade Mountant with DAPI | Invitrogen (Waltham, MA) | Cat# P36966 |
| PVDF Western Blotting Membrane | Roche (Indianapolis, IN) | Cat# 03010040001 |
| Rapamycin | Enzo Life science (Farmingdale, NY) | Cat# BML-A275-0005 |
| Recombinant Human IFN-γ | Peprotech (Rocky Hill, NJ) | Cat# 300-02 |
| Recombinant Murine IFN-γ | Peprotech (Rocky Hill, NJ) | Cat# 315-05 |
| Rhod-2, AM, cell permeant | Invitrogen (Waltham, MA) | Cat# R1245MP |
| RIPA Buffer | Sigma (St. Louis, MO) | Cat# R0278-50ml |

| | | |
|---|---|---|
| Sodium phenylbutyrate (4PBA) | Sigma (St. Louis, MO) | Cat# SML0309 |
| SuperSignal™ West Pico Plus ECL substrate | Thermo Fisher (Waltham, MA) | Cat# 34577 |
| Tetramethylrhodamine, Ethyl Ester, Perchlorate (TMRE) | Thermo Fisher (Waltham, MA) | Cat# T669 |
| Thiazolyl Blue Tetrazolium Bromide (MTT) | Sigma (St. Louis, MO) | Cat# M2128-500MG |
| TNF-α inhibitor | Enzo Life science (Farmingdale, NY) | Cat# ENZ-CHM119-0001 |
| Trizol | Thermo Fisher (Waltham, MA) | Cat# 155596018 |
| ZVAD-FMK | Enzo Life science (Farmingdale, NY) | Cat# ALX-260-020-M001 |
| **Critical commercial kits and assays** | | |
| ATPlite Luminescence Assay System | PerkinElmer (Waltham, MA) | Cat# 6016941 |
| Comet Assay | Trevigen (Gaithersburg, MD) | Cat# 4250-050-K |
| Cytotoxicity Detection Kit PLUS (LDH) | Sigma (St. Louis, MO) | Cat# 4744926001 |
| Direct-zol™ RNA MiniPrep | Zymo Research (Irvine, CA) | Cat# R2052 |
| EnzChek™ Caspase-3 Assay Kit #2 | Thermo Fisher (Waltham, MA) | Cat# E13184 |
| Fast SYBR™ Green Master Mix | Thermo Fisher (Waltham, MA) | Cat# 4385612 |
| Griess Reagent System | Promega (Madison, WI) | Cat# G2930 |
| High-Capacity cDNA Reverse Transcription Kit | Thermo Fisher (Waltham, MA) | Cat# 4368813 |
| Human IL-1β ELISA | BioLegend (San Diego, CA) | Cat# 437004 |
| Human IL-6 DuoSet ELISA | R&D (Minneapolis, MN) | Cat# DY206-05 |
| Human TNF-α DuoSet ELISA | R&D (Minneapolis, MN) | Cat# DY210-05 |
| Mouse anti-dsDNA IgG-specific ELISA Kit | Alpha Diagnostic International (San Antonio, TX) | Cat# 5120 |
| Mouse IL-6 DuoSet ELISA | R&D (Minneapolis, MN) | Cat# Dy-406-05 |
| Mouse TNF-α DuoSet ELISA | R&D (Minneapolis, MN) | Cat# Dy-410-05 |
| RC DC™ Protein Assay Kit | Bio-Rad (Hercules, CA) | Cat# 5000120 |
| RNeasy Plus Mini kit | Qiagen (Germantown, MD) | Cat# 74134 |
| TURBO DNA-free™ Kit | Invitrogen (Waltham, MA) | Cat# AM1907 |
| **Software and Algorithms** | | |
| Biorender | https://biorender.com/ | |
| CFX Maestro | Bio-Rad (Hercules, CA) | Version 2.3 (5.3.022.1030) |
| DAVID bioinformatics database | https://david.ncifcrf.gov/ | Version 6.8 [1] |
| Fiji Software | https://imagej.net/Fiji | Version 1.53c |
| Graph Pad Prism | https://www.graphpad.com/ | Version 8.0 |
| Heatmapper | http://www.heatmapper.ca | [2] |
| ImageJ | https://imagej.nih.gov/ij/ | 1.52a |
| iPathwayGuide | https://advaitabio.com/ipathwayguide/ | Release January 31, 2020 [3] |
| MGI database | http://www.Informatics.jax.org/index.shtml | |
| **Deposited data** | | |
| RNA-seq data | This paper and [4] | GEO accession numbers: GSE173877 GSE159821 |

| **Other** | | |
|---|---|---|
| BX41 Phase Contrast & Darkfield Microscope | Olympus | BX41 |
| Nikon E800 Epifluorescence and Brightfield microscope | Nikon | E800 |
| Real-Time PCR System | Bio-Rad (Hercules, CA) | CFX384 Touch |
| Real-Time PCR System | Thermo Fisher (Waltham, MA) | StepOnePlus |

| **List of primer sequences used for qRT-PCR analysis** | | |
|---|---|---|
| **Gene Mouse** | **Forward** | **Reverse** |
| *Apobec3* | CAGCCATCGCAAATGCTATTC | ATTTCAGCGTGGATGTTGTCC |
| *Cxcl10* | GGATGGCTGTCCTAGCTCTG | TGAGCTAGGGAGGACAAGGA |
| *Dnase1l3* | TCTCGACTTGGAAGAAACACG | GTCTCCATCCTGATAGTCATGGT |
| *Grp78* | ACTTGGGGACCACCTATTCCT | ATCGCCAATCAGACGCTCC |
| *Hprt* | GCCCCAAAATGGTTAAGGTT | TTGCGCTCATCTTAGGCTTT |
| *Ifi44* | GGAGGATTTGCCTTTGAACA | TGGGTTAAAGGTCAGGGCTA |
| *Ifi44L* | ACAGGCTCATGAACCATCCA | TCTCGAGAACTCATGCTCCA |
| *Ifitm1* | GGAGCAGCAAGAGGTGGTTG | GATGTTCAGGCACTTGGCGG |
| *Il-1b* | CAGGCAGGCAGTATCACTCA | TGTCCTCATCCTGGAAGGTC |
| *Irf-7* | TGCTGTTTGGAGACTGGCTAT | TCCAAGCTCCCGGCTAAGT |
| *Mx1-SNE1* | CGGTTGTTTACCAAACTGCG | TTCCAGGGCTTTGACTCGC |
| *Mxi1* | GGCACACAACACTCGGTTTG | CCATTCGTATCCGCTCCATCT |
| *Nos2* | CACCTTGGAGTTCACCCAGT | ACCACTCGTACTTGGGATGC |
| *Oas1a* | GCCTGATCCCAGAATCTATGC | GAGCAACTCTAGGGCGTACTG |
| *Polr2f* | GACAACGAGGACAATTTCGACG | GGAGAATCTCGACATTTTCCTGG |
| *Prkr* | TGCCGTGGTTTTCCTTTAAC | CAGGCCCAGCAATTAACAAT |
| *Rnaseh2a* | GGATAGAGGTGACAGTCAAGGC | CCTGAGCCATAATCGGAGTCCA |
| *Tnfa* | CTGGGACAGTGACCTGGACT | CTCCCTTTGCAGAACTCAGG |
